# Leveraging Twin Information for Inference of Gene Regulatory Networks

**DOI:** 10.64898/2026.02.22.707230

**Authors:** Keerthana M. Arun, Yuval Scher, Yuhan D. Zhang, Ida Büschel, Benjamin Kuznets-Speck, Carsten Marr, Yogesh Goyal

**Affiliations:** Department of Cell and Developmental Biology, Feinberg School of Medicine, Northwestern University, Chicago IL, USA; Center for Synthetic Biology, Northwestern University, Chicago IL, USA; Robert H. Lurie Comprehensive Cancer Center, Northwestern University, Feinberg School of Medicine, Chicago IL, USA; NSF-Simons National Institute for Theory and Mathematics in Biology, Chicago IL, USA; Institute of AI for Health, Computational Health Center, Helmholtz Munich, Neuherberg, Germany; Chan-Zuckerberg Biohub Chicago, Chicago, IL, USA

## Abstract

Gene regulatory networks (GRNs) underlie maintenance of cellular phenotypes and responses to stimuli. Modern single-cell profiling methods offer high-throughput datasets to infer GRNs en masse but do not capture dynamic information. Cell-state heterogeneity further confounds correlation-based inference approaches. We addressed these challenges with TwINFER, a conceptual framework leveraging information from recently divided sister cells, or “twins,” identifiable via recently developed barcoding techniques. We show that twin information discriminates regulatory from non-regulatory correlations and resolves interaction direction and type (activation/repression). We performed a diverse set of simulations, covering common network motifs and large-scale networks, where TwINFER outperforms state-of-the-art inference capabilities. Crucially, TwINFER resolved the commonly observed false positives in fan-out and feed-forward loop motifs where most methods perform poorly. Lastly, we applied TwINFER to a lineage-barcoded hematopoiesis dataset which refined the network inference, flagged multi-state genes, and determined causal relations. Our work exploits cellular twins as untapped information, readily complementing existing inference approaches.

---

Gene activity at the single-cell level is dynamic and variable, leading to molecular heterogeneity even within isogenic cell populations [1–11]. This reflects the underlying stochasticity of transcription dynamics [7–11]— mRNAs often appear in low numbers in the cell, and variations in their counts can be phenotypically consequential [12, 13]. Variability is further amplified because mRNA production occurs predominantly in “bursts” rather than at a constant rate [5–11, 14–20], where each gene transitions stochastically between an inactive state and a short-lived active state.

Typically, transcriptomic states are maintained in cells, with persistent and robust gene co-expression beyond intrinsic noise (illustrated in Figure 1a) [21–23]. Gene expression is regulated by transcription factors (TFs), which modulate the duration and size of transcriptional bursts [10, 20, 23–28]. In turn, TFs themselves are gene products, wherein their production can be regulated by other TFs, forming a gene regulatory network (GRN). Such a network could potentially involve thousands of genes. For instance, the human genome has more than 20, 000 protein-coding genes [21, 29, 30], about 10% of which encode TFs [31, 32]. To mitigate the overwhelming size and complexity of GRNs, studies have identified common network motifs and modules, distinguishable by functional and developmental constraints and operating on different dynamic timescales [33–39].

**FIG. 1.**
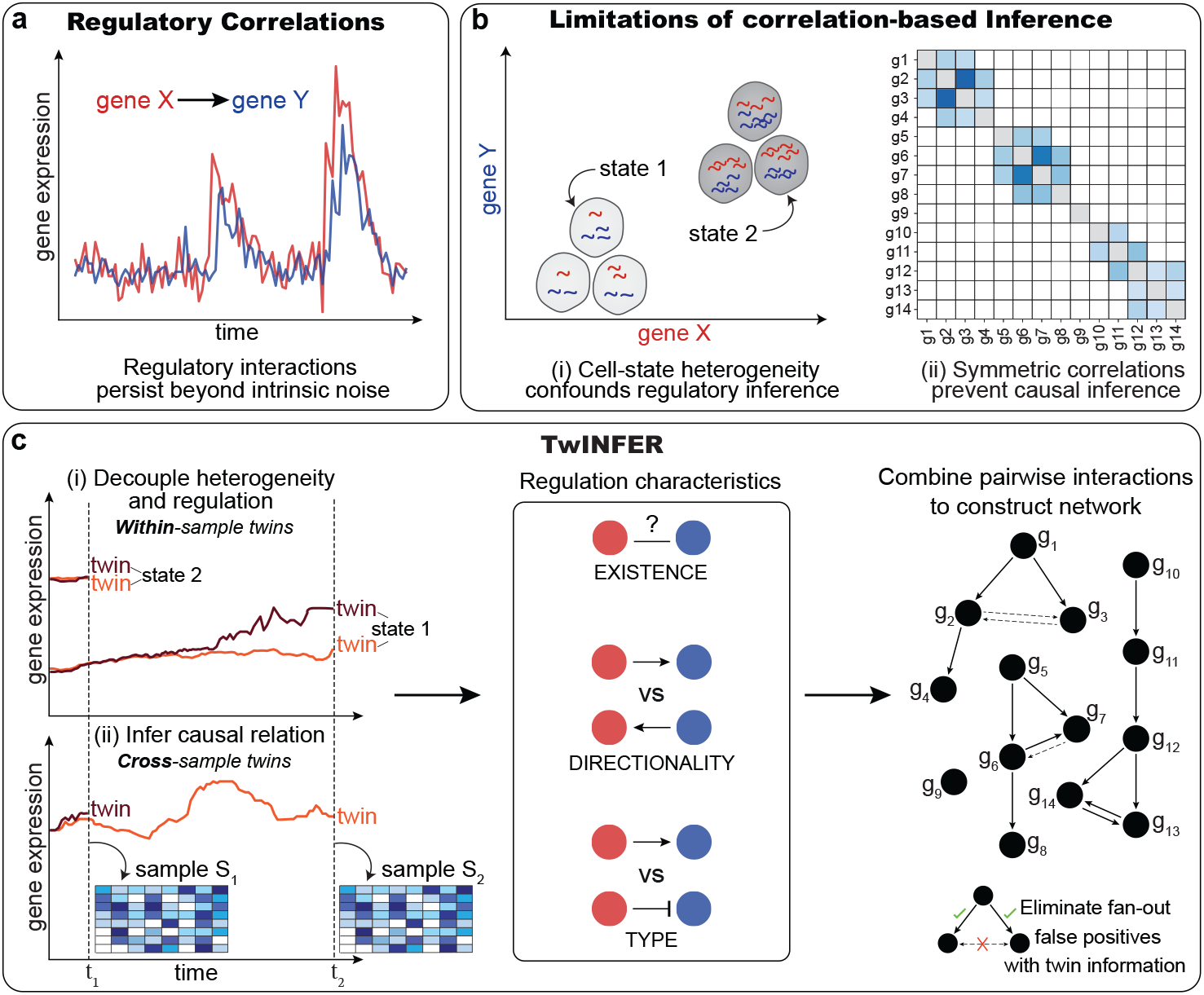
Inferring gene regulatory networks from twin information. **(a)** Gene regulation results in gene co-expression persisting beyond the intrinsic noise. **(b)** We identify two main limitations of correlation-based inference methods: (i) Correlations can arise between genes due to cell-state heterogeneity, even without regulation. (ii) In a GRN, the gene co-expression matrix is symmetric and so causal relations are lost. **(c)** TwINFER uses within- and cross-sample twin correlations measured at two different time points to infer the existence, direction, and type (activation/repression) of regulation. Pairwise interactions can then be combined to infer the gene regulatory network. Twin information can be further used to eliminate false positives in fan-out and feed-forward loop.

As GRNs play a central role in a myriad of cellular processes, inferring their architecture is of paramount importance [35–37, 39–48]. A plethora of computational methods have been devised to leverage regulatory correlations in the transcriptome to reconstruct the underlying GRN, ranging from data-driven methods (e.g., correlation or mutual information networks) to probabilistic models (e.g., maximum likelihood estimation or Bayesian networks) [46–53].

The advent of single-cell transcriptomics [54, 55] permits observation of genome-wide regulatory interactions on a single-cell level, and prompted the development of new inference methods from the growing body of high-throughput data [56, 57]. Oftentimes, modern frameworks integrate additional information, such as *cis*-regulatory motif analysis [56, 58]. Similarly, single-cell multiomics data can be used [48, 57, 59], building on advances in single-cell technologies such as genetic perturbation screens [60, 61], chromatin accessibility [62–64], and chromatin immunoprecipitation assays [65]. Although promising, these approaches are limited in scalability, multiplexing capability, or cost effectiveness [64], and most inference attempts still rely predominantly on transcriptomics [47]. In turn, working with transcriptomic data alone has limitations.

Like bulk profiling techniques, single-cell sequencing is destructive to cells, making dynamic information from within a single cell unattainable [66–68]. One then resorts to looking for regulatory correlations across different cells in the populations. We identify two main obstacles (Figure 1b): (i) Correlations across an isogenic cell population may arise from other non-regulatory sources, obfuscating the desired regulatory interactions. Markedly, instances of two and three co-existing sub-populations with different transcriptional states are often encountered, with a dynamic timescale much longer than typical regulatory correlations [69, 70]. As an example, imagine a scenario of two genes that are both highly expressed in one state and lowly expressed in the other. Due to the heterogeneity in cell states, these genes will be correlated across the population, confounding inference of regulatory interactions [71]. (ii) The symmetric nature of gene-expression correlations renders inferred networks directionless, and causal relations between regulators and their target genes are lost [46, 53, 57, 72, 73].

Here, we introduce TwINFER (Figure 1c), a framework that addresses both of these obstacles by incorporating twin information: a broad term used to denote any additional information that can be extracted by identifying pairs of sister cells. TwINFER discriminates between regulatory and non-regulatory correlations, and determines causal relations. We demonstrate the power of our framework by testing it on an extensive set of synthetic GRNs, where the ground truth is known. The simulations were based on a detailed interaction model, and the parameters were carefully chosen based on published experimental data (Box 1) [17, 18, 74–80]. To establish that our technique is based on fundamental system-independent ideas, in the Supplemental Material we considered a linear toy model. This linear model is analytically tractable such that we were able to recapitulate our framework with mathematical rigor. We began with the building block of GRNs: a system of two interacting genes. For this paradigmatic set-up, we exhausted all possible combinations for the direction and type (activation/repression) of the interaction. We then tackled larger networks of increasing size and complexity, explicitly addressing combinatorial regulation [81] and motifs that are known to be difficult to infer using correlation-based inference [82]. Relying on the known ground truth of in silico experiments, we showed that TwINFER out-performs the state-of-the-art transcriptomic-based GRN inference method GRNBoost2 [52]. Finally, we implemented TwINFER on lineage-barcoded single-cell RNA-sequencing (scRNA-seq) dataset [83].

Collectively, our study demonstrates how twin information can identify causal interactions, cell-state heterogeneity, and illusive triplet motifs. It does so in both synthetic and real experimental datasets, and is especially suitable for emerging barcoded scRNA-seq datasets across different cell types and contexts [83–88].

## Results

### Overview of TwINFER

What is twin information? A storied encounter Alice has during her adventures in the looking-glass world is with the twin pair Tweedledee and Tweedledum [89]. Their portrayal embodies the world in which they live—they are the spitting image of each other, as if one is the reflection of the other. Although in a quarrelsome manner, they end up complementing each other’s sentences. TwINFER essentially relies on the same notion when analyzing the transcriptomes of recently divided sister cells. That is, we assume that a cell approximates its sister cell’s state well. We refer to a pair of such cells as “twins.”

We exploit two kinds of “twin information” in TwINFER to overcome the two aforementioned obstacles: accounting for cell-state heterogeneity and inferring causal relations. This is illustrated in Figure 1c: (i) Within-sample information, where we measure twin pairs from the same sample at the same time point; (ii) Cross-sample information, where a twin and its counterpart are separated between two samples collected at different time points. Overall, we measure the transcriptomes of two samples, S_1_ or in S_2_, at the corresponding time points *t*_1_ and *t*_2_, such that *t*_1_ *< t*_2_. Note that a pair of within-sample twins can be found either in S_1_ or in S_2_. Importantly, *t*_1_ is chosen such that sample 1 is measured shortly after the mother cells divided. The choice of *t*_2_ is more subtle and will be discussed below.

We began by corroborating our results on the paradigmatic set-up of two interacting genes (Figures 2 and 3). In Box 1 and Methods we elaborate on our modeling, the literature-based choice of biologically relevant parameters, and the simulation algorithm. A sketch of the interaction model is provided in Figure E1.

**FIG. 2.**
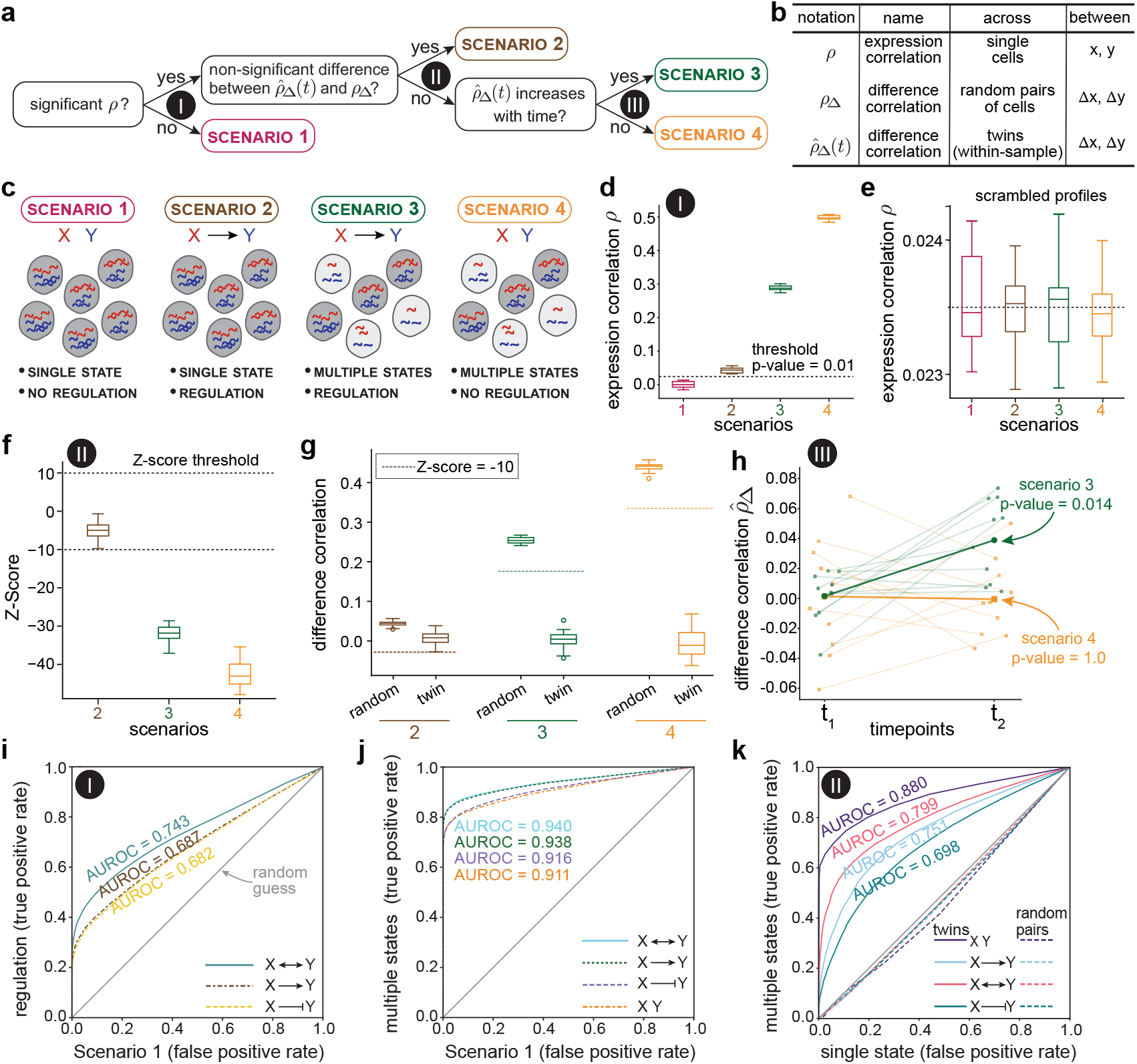
Twin difference correlations account for cell-state heterogeneity. **(a)** An analysis flowchart for discerning whether correlations arise from regulatory interactions, cell-state heterogeneity, or both. The three stages of the flowchart are enumerated with Roman numerals. In case of regulatory correlations, one can proceed to determine the interaction direction (Figure 3). **(b)** A table defining the different types of correlations used to account for cell-state heterogeneity. Here we introduce pair information, with two additional types of correlations. We use a hat 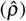 to denote twin correlations. When the correlations are within-sample, we use a ‘Δ’ subscript to denote that the correlations are between the *differences* in gene expressions within the pair, Δ*x* and Δ*y*, rather than the correlations in transcriptional counts, *x* and *y*. **(c)** Schematic of the four scenarios (color-coded). **(d-h)** Parameters chosen: 6, 000 twin pairs; 20 repetitions per box plot; *t*_1_ = 1 *h* and *t*_2_ = 20 *h*; median and default values from Tables E1 and E2 (see Methods and Box 1 for details). Heterogeneity in Scenarios 3 and 4 was modeled with different bursting frequency *k*_on_ for all genes: 0.12 *h*^−1^ and 1.66 *h*^−1^ (see Methods and Figure E7 for details). **(d)** Test for stage I: permutation test over 10, 000 scrambled gene-expression profiles (Figure E3) where one-tailed p-value of 0.01 ensured zero misclassifications. **(e)** Thresholds across four scenarios are indistinguishable and can be used interchangeably. **(f)** Test for stage II: Compute ρ_Δ_ and 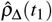, perform permutation test over 10,000 random pairings (Figure E4) and z-test for hypothesis testing on 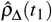. **(g)** With our model and parameters, setting a z-score of 10 ensures zero misclassifications. **(h)** Test for Stage III: this step requires multiple repetitions/sub-sampling of the data for statistical testing (Wilcoxon signed-rank test, −10 repetitions, *p <* 0.05 for Scenario 3 and *p* = 1 for Scenario 4). **(i, j, k)** receiver-operator characteristic curves (ROC) for TwINFER over 25, 000 Latin-hypercube-sampled parameter sets (ranges in Tables E1 and E2 and details in Methods) for regulation **(i)** and multiple-states **(j)**, with area under ROC (AUROC) values provided. **(k)** ROC for the detection of cell-state heterogeneity using stage II of the flowchart for TwINFER and corresponding random pairs.

**FIG. 3.**
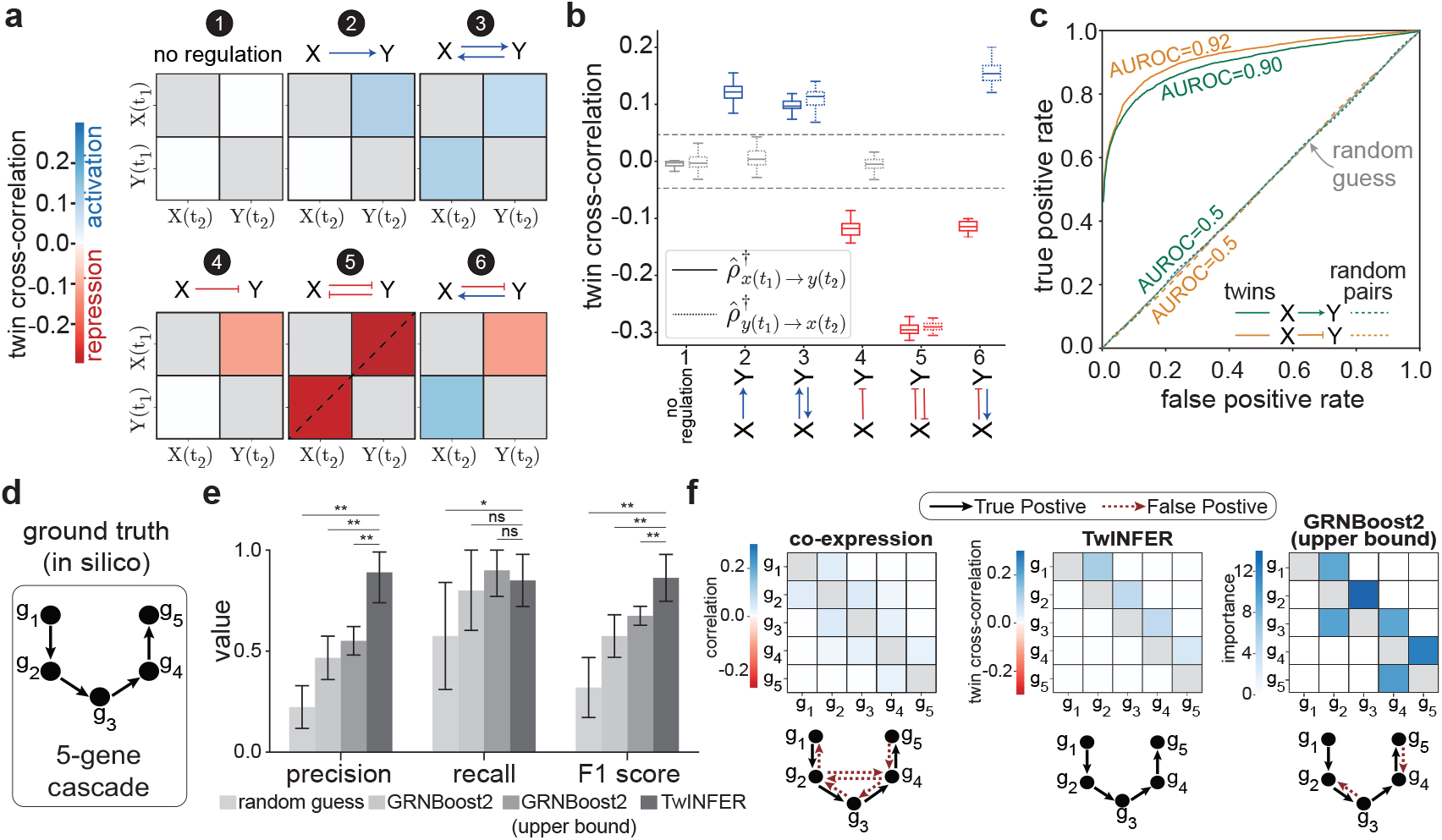
Twin cross-correlations allow inference of causal relations. **(a)** Cross-correlation matrices for five pairwise interaction scenarios between genes X and Y, exhausting all combinations of direction (univs. bidirectional) and type (activation vs. repression), plus a no-interaction control; parameters as in Figure 2. Two cases are irregular: Case 5 (bistability from mutual repression ([94]) and Case 6 (false negative from mixed-sign bidirectional regulation, detectable only via cross-correlations). **(b)** Threshold correlations 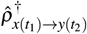 and 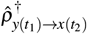 for each case, determined by permutation test over 10,000 random-pairing scrambles (two-tailed *p* = .02; Figure E5b). **(c)** ROC curves for inferring interaction direction using twin vs. random-pair information (AUROC values provided); parameter space scanned over ranges in Tables E1 and E2 using 25,000 Latin-hypercube-sampled sets for activation and repression. **(d)** A linear cascade of five genes simulated as ground truth for evaluating inference. **(e)** Bar plot depicting mean and SD of precision, recall, and F1 score over 10 simulation repeats. TwINFER is compared with random guessing, gene co-expression, GRNBoost2 [52] (two cases: threshold via Gaussian Mixture Model, and an upper bound identified retrospectively for the best threshold; Methods and Figure E8). Scores compared using two-sided paired Wilcoxon test; **, p ≤ 0.01; *, p < 0.05; ns, not significant. **(f)** Representative inferences showing correlation/importance matrices and inferred networks for standard gene co-expression, TwINFER, and GRNBoost2.

### TwINFER reconstructs gene interactions for heterogeneous data

When a cell divides, its molecular content is shared between the resulting twins. Shortly after division, the transcriptomes and proteomes of a twin pair are often very similar [83, 91–93]. Indeed, both sister cells initially belong to essentially the same state and the initial *difference* in the gene expression levels between them is minimal. Formally, consider a gene X and a pair of cells expressing it with time-dependent transcript counts *x*(*t*) and *x*^′^(*t*). We denote their difference Δ*x*(*t*)= *x*(*t*) − *x*^′^ (*t*). At *t*_1_, shortly after the division, if the pair comprises twin cells we have Δ*x*(*t*_1_) ∼ 0. The same holds for a gene Y.

For the case with no regulatory interaction between X and Y, an additional possible source of correlations is the existence of multiple transcriptional states that cells in the heterogeneous population can occupy (or a continuous cell-state heterogeneity due to extrinsic noise [1]). A simple intuitive example would be a population with two states: in one state the expression of both genes is low, and in the other it is high. If we happen to measure a cell with a low X count, it is highly likely that the cell belongs to a low-expression state, and therefore Y counts are low as well, and vice-versa, resulting in correlations. However, for twins Δ*x*(*t*_1_) ∼ Δ*y*(*t*_1_) ∼0, regardless of the initial state of the twins. Hence, Δ*x*(*t*_1_) and Δ*y*(*t*_1_) are uncorrelated across a population of twin pairs. This remains true at a later time point *t*_2_ (given that *t*_2_ is much smaller than the dynamic timescale for state transitions), which we explore systematically in Figure E2a.

Let us now consider the case in which X regulates Y. Still, Δ*x*(*t*_1_) ∼ Δ*y*(*t*_1_) ∼ 0 for twins. However, the regulatory interaction leads to a correlated change in transcript counts throughout the interval [*t*_1_, *t*_2_], and we expect Δ*x*(*t*_2_) and Δ*y*(*t*_2_) to become more correlated as *t*_2_ increases (see Figure E2b). Based on this intuition, we present a flowchart that describes the analysis pipeline to overcome the obstacle of cell-state heterogeneity (Figure 2a). Figure 2b summarizes the definitions and notations of all the different correlations that appear in the flowchart. The four different outcomes of the flowchart are: 1. No regulation and a single state; 2. Positive regulation and a single state; 3. Positive regulation and two states; 4. No regulation and two states (summarized in Figure 2c). Note that we use Spearman’s rank correlation coefficient to measure correlations. We outline the flowchart stepwise below.

We began by calculating *ρ*, the ordinary gene-expression correlation across a population of single cells (merging the data from samples S_1_ or in S_2_, as *ρ* is time-independent, see Methods). Following the seminal algorithm ARACNE [50], we set a statistical threshold below which the null hypothesis of mutually independent genes cannot be ruled out. We performed a permutation test with 10,000 shuffled gene expression profiles (Figure E3). If *ρ* is below the threshold, we conclude that there is no significant correlation. That is, there is a single state without regulation. A value of *ρ* above the threshold is a necessary condition to determine the existence of regulation, but not a sufficient condition, as cell-state heterogeneity can also result in correlations. To determine whether there is cell-state heterogeneity, we compute the correlation between Δ*x*(*t*_1_) and Δ*y*(*t*_1_) across within-sample pairs of twins, denoted by 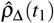. We then perform a different type of permutation test: we compute the time-independent correlations between Δ*x* and Δ*y*, denoted by *ρ*_Δ_, across 10,000 random pairings of cells. If the difference between *ρ*_Δ_ and 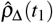 is below the statistical threshold (Figure E4), we conclude that there is a single state and therefore the source of gene correlation is due to regulatory interactions.

#### Box 1.

Model Description and Parameters

**Model Description**

The transcriptional bursting of unregulated genes is modeled as a telegraph process. Each gene transitions between two states, ‘on’ and ‘off’. Transcription occurs only in the on state, and is followed by transcription factor (TF) synthesis. TFs and mRNAs are degraded at constant rates.

The chemical reactions and propensities are described in the table below:

| Reaction | Propensity |
| --- | --- |
| $x_{\text{off}} \rightarrow x_{\text{on}}$ | $k_{\text{on}} \cdot x_{\text{off}}$ |
| $x_{\text{on}} \rightarrow x_{\text{off}}$ | $k_{\text{off}} \cdot x_{\text{on}}$ |
| $x_{\text{on}} \rightarrow x_{\text{on}} + \text{mRNA}_x$ | $k_{\text{prod}}^{\text{mRNA}} \cdot x_{\text{on}}$ |
| $\text{mRNA}_x \rightarrow \text{mRNA}_x + \text{TF}_x$ | $k_{\text{prod}}^{\text{TF}} \cdot \text{mRNA}_x$ |
| $\text{mRNA}_x \rightarrow \emptyset$ | $k_{\text{deg}}^{\text{mRNA}} \cdot \text{mRNA}_x$ |
| $\text{TF}_x \rightarrow \emptyset$ | $k_{\text{deg}}^{\text{TF}} \cdot \text{TF}_x$ |

where *x*_on_, *x*_off_ ∈ [0, 1], and *x*_on_ +*x*_off_ = 1.

Regulatory interactions are manifested through the modulation of the bursting process by TFs. For example, let gene X be a regulator of gene Y. The TF synthesized by gene X modifies the rate of bursting of gene Y. Note that the bursting of gene Y ceases to be a simple telegraph process, as the Markov property is lost [90]. The burst frequency changes from *k*_on_ in the absence of regulation to 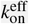 under regulation according to

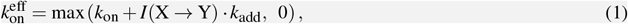

where *I*(*X* →*Y*) is an indicator function that returns the value 1 if X is an activator and −1 if X is a repressor. The strength of regulation is determined by *k*_add_, and it increases with the levels of the TF_*x*_, until it reaches a plateau. This behavior is captured by the Hill function

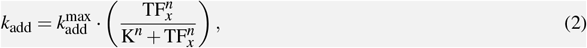

where 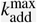 is the maximum possible change, *n* is the Hill coefficient, and K is the Hill constant.

**Parameters - Transcriptional Bursting**

Gene expression in the absence of regulation is described by 7 model parameters, as defined in the table below along with corresponding ranges based on literature values.

| Parameter | Description | Range | Median | Units | References |
| --- | --- | --- | --- | --- | --- |
| $k_{\text{on}}$ | Rate of burst initiation | 0.01 – 3 | 0.66 | $h^{-1}$ | [17, 18, 74] |
| $k_{\text{off}}$ | Rate of burst inactivation | 0.06 – 55 | 8.6 | $h^{-1}$ | [17, 18] |
| $\pi_{\text{on}}$ | Bursting fraction | 0.002 – 0.4 | 0.07 | – | [17, 18] |
| $k_{\text{prod}}^{\text{mRNA}}$ | Transcription rate | 0.2 – 60 | 2 | $\text{mRNA} \cdot h^{-1}$ | [17, 18, 75, 79] |
| $k_{\text{prod}}^{\text{TF}}$ | Translation rate | 19 – 2700 | 560 | $\text{TF} \cdot \text{mRNA}^{-1} \cdot h^{-1}$ | [75, 79] |
| $\tau_{1/2}^{\text{mRNA}}$ | mRNA half life ($1/k_{\text{deg}}^{\text{mRNA}}$) | 0.6 – 17 | 4 | $h$ | [18, 76–79] |
| $\tau_{1/2}^{\text{TF}}$ | Protein half life | 7 – 200 | 45 | $h$ | [79, 80] |

where *τ*_1*/*2_ = 1*/k*_deg_ and *π*_on_ = *k*_on_*/* (*k*_on_ +*k*_off_).

**Parameters - Regulatory Interaction**

Each regulatory interaction requires three additional parameters, where 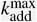 takes different values for positive (+) or negative (−) interactions. The range for possible 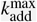 values is a function of *k*_on_ (linear scaling).

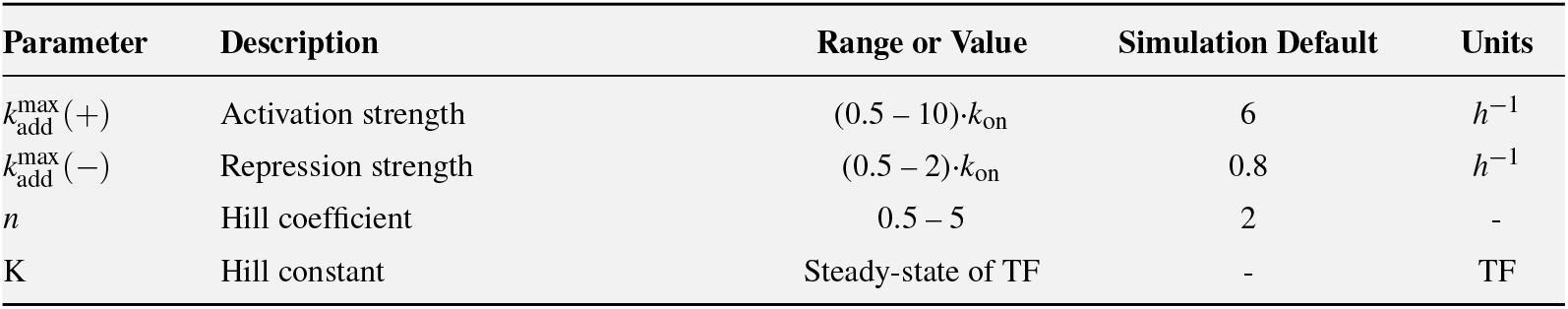

If the difference between *ρ*_Δ_ and 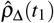 is above the threshold, we conclude that there is cell-state heterogeneity. However, we cannot yet conclude whether there is regulation or not, as one must consider the possibility of heterogeneity *and* regulation. In the last step, we compare 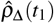 and 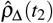. A significant increase during this time implies the existence of heterogeneity and regulation.

We challenged our method with simulations of the four scenarios (see Methods for more details on the modeling, parameter choice, and simulation algorithm). By following the flowchart (Figure 2a), we were able to infer them correctly. Importantly, stages I-II, both relying on permutation tests, require only a single repetition of the experiment (we simulated 6,000 mother cells per experiment). We demonstrated the statistical power and repeatability of the results in Figure 2d-h. The panels are divided into three groups (d-e, f-g, and h), each representing a test of the possible scenarios in accordance with the corresponding stage of the flowchart. As expected, only Scenario 1 had insignificant gene co-expression (Figure 2d), when we set a threshold p-value in the shuffle test (Figure 2e shows that the scrambled profiles of all scenarios are essentially indistinguishable and can be used interchangeably). Similarly, of the remaining three scenarios in Figure 2f-g, only Scenario 2 had an insignificant difference between random-pair and twin correlations. Lastly, we distinguished between Scenarios 3 and 4 in Figure 2h.

To further challenge TwINFER and better define its inference capabilities and limitations, we scanned the parameter space restricted by the ranges in Tables E1 and E2, testing 25, 000 Latin-hypercube-sampled parameter sets for each of the scenarios (Figure 2i-k; see Methods). We scanned the cases of repression and bidirectional interactions as well. We plotted the receiver-operator characteristic (ROC) curves for the detection of regulation (Figure 2i) and multiple-states (Figure 2j). False positives occur when Scenario 1 was simulated, but was inferred as one of the other scenarios, and where the test is performed according to stage I of the flowchart. We next plotted the ROC curves for the detection of cell-state heterogeneity using stage II of the flowchart (Figure 2k). Overall, we obtained a high area under the receiver operator curve (AUROC) values (range = 0.68-0.94; median = 0.80), demonstrating that TwINFER performs well across a wide range of parameter values.

### Inferring causal relations

In cases where the source of the correlation is established solely as regulatory, we proceeded to infer causal relations. We accomplished this by turning to cross-sample twin information, where a twin and its counterpart are separated between the two samples S_1_ and S_2_, measured at time points *t*_1_ and *t*_2_, respectively.

Recall that at *t*_1_, shortly after division, a cell approximates its twin very well. As a result, initially, their transcriptomes follow similar trajectories in time. Due to the stochastic nature of gene expression, the trajectories will gradually diverge (Figure 1c). Eventually, the twins will dephase completely—given enough time, it becomes impossible to determine whether a pair of cells started as twins or not on the basis of the transcriptome alone. When setting biologically relevant parameters in our model, we observe that the twins’ trajectories retain high similarity for many hours. Thus, as long as *t*_2_ falls within this time frame (Figure E5), the state of a cell in S_2_ approximates well the state that its counterpart in S_1_ would have reached during the time interval [*t*_1_, *t*_2_] if not interrupted by a destructive transcriptome measurement. Each sample can only be measured once, and dynamic information is unavailable, precluding inference of causal relations. TwINFER exploits the similarity between the trajectories of twins separated between the two samples, treating them as a single trajectory measured at two different time points. In this manner, we obtain sufficient dynamic information to resolve causal relations.

We used the cross-sample “cross-correlations” between the transcript counts in the two samples: For the direction X→Y, we computed the cross-sample correlation between *x*(*t*_1_) and *y*(*t*_2_), denoted here 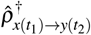. For the reverse direction Y → X we computed the cross-sample correlation between (*t*_1_) and *x*(*t*_2_), denoted here 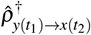. We reasoned that if the interaction is unidirectional, only one of these cross-correlations will be significant, revealing the causal relation. If the interaction is bidirectional, both cross-correlations will be significant. Furthermore, the sign of the cross-correlation determines the type of the interaction: Positive and negative signs correspond to activation and repression, respectively.

We next challenged TwINFER with simulations of all possible types of pairwise interactions (Figure 3a), considering unidirectional and bidirectional interactions, and allowing both activation and repression. See Methods for more details on the modeling, parameter choice, and simulation algorithm. Markedly, the cross-correlation matrices were no longer symmetric, and the causal relations were correctly inferred in all cases (Figure 3b, 20 repetitions per case). ROC analysis across the parameter space (Tables E1 and E2) yielded AUROC ∼0.9 for both activation and repression (Figure 3c). As a sanity check, when random pairs were used instead of twins, we obtained AUROC scores of ∼ 0.5 (Figure 3c), equivalent to random guessing.

Throughout Figure 3, Sample 1 and Sample 2 were measured at *t*_1_ = 1 and *t*_2_ = 20 hours after division, respectively. To check for robustness of TwINFER under different measurement times, we systematically scanned all possible pairs of *t*_1_ and *t*_2_, computing the cross-correlations in the direction of the simulated interaction (Figure E5a). Briefly, for our median parameters, correct inference was obtained when Sample 1 is measured within ∼5 hours after division. Sample 2 had a much wider window, peaking at ∼15 −35 hours, for measuring *t*_2_. Furthermore, we provide an example of the permutation test to determine the significance of the directional interaction (Figure E5b).

Lastly, we considered a linear toy model of gene interactions representing a linearization about a local transcriptional attractor (see Supplemental Material). This model is analytically tractable, so that we were able to provide a mathematical proof for the ideas that were postulated in our treatment of the non-linear model. The observation that the results of the analytically solvable model carry over to our far more elaborate model suggests that the underlying ideas presented here may be universal.

### On inference and simulation of larger networks

Next, we turned our attention to networks that comprise more than two genes. When inferring larger networks, we applied the pairwise pipeline to each gene pair independently and consolidated the inferred interactions to reconstruct the full network.

To simulate cascades and cascade-derivatives with the Hill constants K set to steady-state regulator concentrations, we performed a hierarchical series of simulations of cascades increasing in size. When the regulators are themselves regulated by upstream genes, the steady state is governed by Eq. (E10), which must be evaluated numerically by simulating the cascade up to that gene, see Methods (note that if we were to continue using the time-efficient analytical Eq. (E8), the defined K would be off by a few percents from the steady-state concentration, which was found to be inconsequential (Figure E6c)). Here, for completeness, we used the hierarchical procedure when simulating the 5-gene cascade, and we show its successful reconstruction in Figure 3d-e.

To quantify the performance of TwINFER we repeated the simulation of the 5-gene cascade 10 times and computed the mean and standard deviation of its precision, recall and F1 score. In Figure 3d we compare this result with: (i) Random guessing; (ii) Gene co-expression, which does not allow for determination of the direction, and therefore will have at least 4 false positives via prediction of all reverse directions; (iii) GRNBoost2 [52], which gives an importance score to all potential interactions, and where we choose the importance threshold based on a Gaussian Mixture Model (Methods and Figure E8); (iv) The upper bound for GRNBoost, i.e., retrospectively choosing the best performing threshold (highest F1 score) in order to define a bound for GRNBoost2 performance. To illustrate a representative result, we plotted the correlation/importance matrices and their corresponding inferred networks (Figure 3e).

We chose GRNBoost2 as a representative of state-of-the-art capabilities for GRN inference. GRNBoost2 is based on the GENIE3 algorithm [51], winner of the DREAM5 GRN inference challenge [52]. By integrating twin information, the average precision of TwINFER is two-fold higher than that of GRNBoost2 (Figure 3d). In fact, TwINFER’s mean precision is almost 1.

Lastly, in larger networks, a gene may be simultaneously regulated by multiple TFs. We therefore extended our interaction model to account for such a combinatorial regulation as done in Ref. [81]. For example, consider two activators of the same *cis*-regulatory region. We have four possible states, corresponding to the combinations of concomitantly bound TFs. In Methods, we generalized the interaction’s Hill function to account for their joint effect and retrieve paradigmatic cases. In what follows, we employed the additive case in which the different interactions are independent and the overall regulatory effect is the sum of pairwise interactions.

### The fan-out motif as a case study

We expanded the two-gene system by adding a gene Z that can pairwise interact with both X and Y. For each edge in the triplet there are four options: no interaction, unidirectional interaction in the forward direction, unidirectional interaction in the reverse direction, and bidirectional interaction. Overall, the addition of a single gene increases the number of possible configurations from 4 to 64 (even without considering repression).

Some of these configurations are known to be difficult to infer. Among these, the fan-out motif (Figure 4a) is especially notorious [82], as it is also an ubiquitous motif in real biological contexts, along with the feed-forward loop (Figure 4a) [33, 72]. These two important motifs are indistinguishable based on gene co-expression and many inference methods fail to reconstruct them [82].

**FIG. 4.**
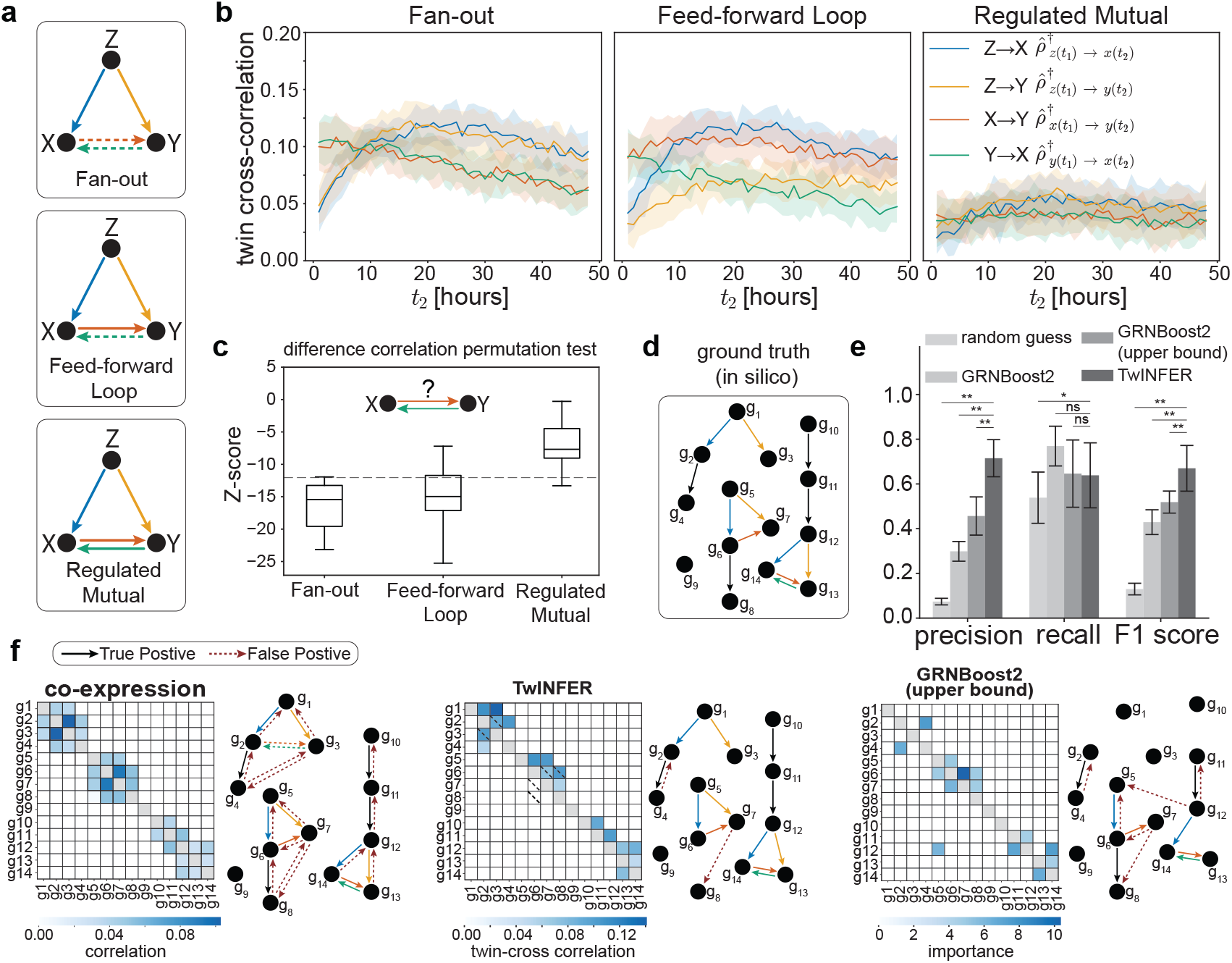
Twin cross-correlations distinguish between triplet motifs. **(a)** Three triplet motifs with dashed lines denote false positives from correlation-based inference. **(b)** Corresponding twin cross-correlations vs. *t*_2_; *t*_1_ = 1 *h*. Divergence between 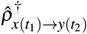 and 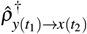 indicates directional asymmetry. **(c)** Box plots of z-scores from the twin difference correlation permutation test for the X-Y interaction across the triplet motifs (20 repetitions). A z-score threshold of −12 separates fan-out and feed-forward loop from regulated mutual. **(d)** Ground truth network for evaluating inference. **(e)** Bar plot depicting mean and SD of precision, recall, and F1 score over 10 simulation repeats on the network in **(d)** for (i) random guessing, (ii) gene co-expression, (iii) TwINFER, and (iv) GRNBoost2 [52] (two cases: threshold via Gaussian Mixture Model, and an upper bound identified retrospectively for the best threshold; Methods and Figure E8). Scores compared using two-sided paired Wilcoxon test; **, p ≤ 0.01; *, p < 0.05; ns, not significant. **(f)** Representative inferences showing correlation/importance matrices and inferred networks for standard gene co-expression, TwINFER, and GRNBoost2. Default parameters from Tables E1 and E2 (and Box 1) were used in **(a-f)**, and combinatorial interactions were additive (see Methods).

When the ground truth motif is fan-out, correlation-based inference methods often infer it as the regulated mutual motif (Figure 4a). In fact, the fan-out motif represents a classical problem in inference, where gene Z acts as a confounding factor. Moreover, the feed-forward loop motif can also be incorrectly inferred as a regulated mutual. Therefore, when the result of a correlation-based inference method is a regulated mutual, it is likely that at least one of the inferred interactions X → Y and X ← Y is a false positive. Given the ubiquity of fan-out motifs in GRNs, we faced the question: Upon inferring a regulated mutual motif, can we leverage twin information to assess the likelihood of this edge being a false positive?

Here again, we utilized both within- and cross-sample twin information. First, we show that twin cross-correlations 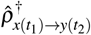 and 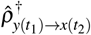 are significantly different in the feed-forward loop case, reflecting its asymmetry and distinguishing it from the two other motifs (Figure 4b).

The key to discerning between fan-out and regulated mutual is more subtle. Note that the confounding variable Z is inherited from the mother cells, such that twins share the same initial concentration of Z. However, across the population, the concentration of Z varies (drawn from its steady-state distribution). In essence, despite being part of the network, and from the perspective of the interaction between X and Y, Z functions as extrinsic noise. This means that the same technique used to detect cell-state heterogeneity can also detect the presence of the confounding variable. In other words, the interaction between X and Y in the fan-out case will be flagged as heterogeneous, with a low (negative) z-score in the difference correlation permutation test (Figure 4c).

In contrast, the impact of Z on X and Y is attenuated if they are part of an embedded fan-in motif, namely if they are concomitantly regulated by two TFs (e.g., Y is regulated by both X and Z in the feed-forward loop). The regulated mutual contains two embedded fan-in motifs, resulting in all its interactions exhibiting substantially low correlations (Figure 4b). This observation was further supported by our analysis on all sub-motifs of regulated mutual, where only fan-in results in a comparable attenuation (Figure E9). Moreover, the embedded fanin within the regulated mutual motif suppresses the difference correlations *ρ*_Δ_ between X and Y. These analyses enabled us to distinguish regulated mutual from fanout using the heterogeneity test (Figure 4c). (Although fan-in correlations are attenuated in our linear model as well, the two models respond qualitatively differently to the mutual activation of X and Y. In the linear model, cross-correlations are sufficient to distinguish between the three motifs (see Supplemental Material).)

### Putting it all together in a complex network

We next asked whether TwINFER is powered to infer a relatively large, complex network (Figure 4d). TwINFER refined the prediction further by inferring causal relations, as measured by precision, recall, and F1 scores of 10 simulation repeats (Figure 4e). As in the previous subsection, we compared between TwINFER’s performance, GRNBoost2 thresholded with the Gaussian Mixture Model (Figure E8), and GRNBoost2’s upper bound (Methods). TwINFER’s precision is about two-fold higher than that of GRNBoost2. Among all methods, only TwINFER differentiates between sources of correlations. We chose to evaluate TwINFER stringently by counting as false positives the few cases where TwINFER correctly inferred the presence of a correlation but misidentified its source. Consequently, TwINFER’s recall is slightly reduced.

The simulated network comprises the three triplets discussed in the previous section. For a representative repeat, we show the inferred matrices and corresponding GRNs (Figure 4f). The gene co-expression matrix, when naively interpreted, resulted in the common false positives for fan-out and feed-forward loop, while TwINFER correctly identifies them (possible false positives are flagged with diagonal dashed lines). While the errors of the correlation-based inference methods are contained within local elements of the networks, GRNBoost2 infers spurious connections between isolated sub-elements of the network (Figure 4f, rightmost panel).

### Application of TwINFER on experimental data

We next turned our attention to testing TwINFER on experimental data. Recent developments in single-cell DNA barcoding methodologies [83–88] have enabled tagging individual cells uniquely and following the fates of clonally related cells. As such, uniquely barcoded mother cells divide and pass on their barcode to their off-springs (Figure 5a).

**FIG. 5.**
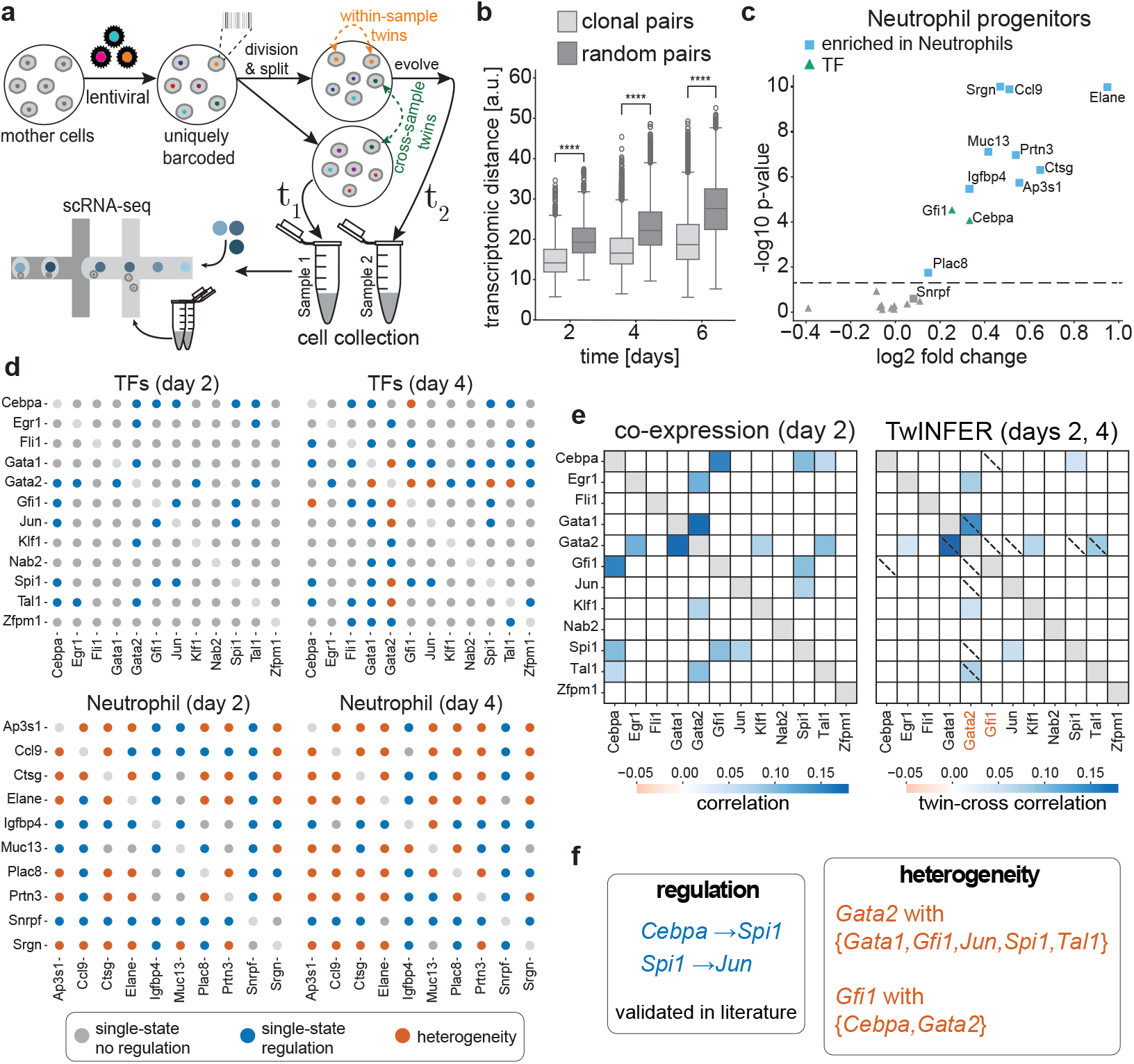
TwINFER captures regulatory interactions and confounding heterogeneity in experimental data. **(a)** Schematic of the experimental design: mother cells receive unique barcodes via lentiviral transduction; daughter cells inherit barcodes, forming twin pairs that are split and measured at *t*_1_ and *t*_2_. **(b)** Box plots of transcriptomic distance between clonal and random pairs at days 2, 4, and 6 in the LSK LARRY dataset [83] (we used singletCode [85] for preprocessing; see Methods). Distances compared by two-sided Mann-Whitney U test; ****, *p ≤* 10^−4^. **(c)** Volcano plot (corresponding to Figure 2G of Ref. [83], reanalyzed with our approach) identifying genes enriched in neutrophil progenitors relative to the general population at day 2. Two gene groups are tested: neutrophil-enriched genes reported in [83] (rectangles) and TFs driving murine myeloid differentiation [95] (triangles). **(d)** Dot matrices for TF and neutrophil gene groups at days 2 and 4, with dot color denoting the inferred scenario per gene pair (Figure 2; Scenarios 3 and 4 merged into a single heterogeneous multi-state category). **(e)** Gene co-expression matrix (day 2, left) and TwINFER cross-correlation matrix (days 2 and 4, right) for gene pairs, after we filtered for pairs with stable correlations (Methods, Figure E11). **(f)** Summary of TwINFER-identified heterogeneity (*Gata2* with *Gata1, Gfi1, Jun, Spi1, Tal1*; *Gfi1* with *Cebpa, Gata2*) and regulatory interactions (*Cebpa*→*Spi1, Spi1*→*Jun*), both interactions are validated in the literature [97–99].

We used the publicly available Lin^−^Sca1^+^Kit^+^ (LSK) LARRY dataset of Ref. [83], which contains scRNA-seq data from LSK mouse hematopoietic progenitors. These were cultured in a medium designed to support pan-myeloid differentiation. The barcoded mother cells were cultured for 2 days, at which point their daughter cells were split into three samples, measured at three different time points: after 2 days (immediately after splitting), 4 days, and 6 days. These measurement times were longer than the timescales for TwINFER discussed above and in Figure E5. During the experiment, cells divided multiple times, thus the twin pairs used in the analysis were clonal pairs rather than strict sister cells. Nonetheless, clonal pairs were significantly more similar to each other based on transcriptomic distance, and this property was broadly retained throughout the experiment (Figure 5b).

Figure 5c follows Figure 2G of Ref. [83], but is repeated with our preprocessing and analysis protocols (see Methods). We linked undifferentiated progenitors to their most likely fate using barcode-matched clonal data from days 4 and 6. Cells were categorized by the most frequent fate among their cross-sample clones. We chose to focus on myeloid differentiation as cells with other fates (lymphoid or pDC) were present in relatively smaller numbers. Within the myeloid lineage, we analyzed neutrophils (Figure 5) and monocytes (Figure E10). The volcano plot identifies genes enriched in neutrophils compared to the general population on day 2. Of 10 genes reported in Ref. [83], 9 were significantly enriched in our data. However, of 12 transcription factors known to drive murine myeloid differentiation [95], only two (*Cebpa* and *Gfi1*) were significantly enriched in neutrophils.

For each group of genes, we summarize the results of applying the flowchart from Figure 2 to all possible pair-wise interactions (Figure 5d). The TF-encoding genes are exclusively homogeneous on day 2, while correlations of highly enriched genes in neutrophils often originate from heterogeneity: approximately half of the interactions are flagged as heterogeneous (orange circles). This confirms the intuition that differentially expressed genes tend to be in multiple transcriptional states. Note that while the identification of differentially expressed genes requires subpopulations categorized by fate, day 2 sample is sufficient for TwINFER to predict heterogeneity. Lastly, we emphasize that differentially expressed genes may show significant correlations, and these were shown here to emanate, at least partially, from heterogeneity of transcriptional states—indeed, one should be cautious when interpreting gene co-expression as evidence for regulation.

We applied TwINFER to the network of TFs encoding genes. As a preliminary step, we identified correlations that remain significant and of the same type throughout the experiment, which we call “stable correlations” (Figure E11). We compared the gene co-expression (day 2) matrix and the twin cross-correlation (between days 2 and 4) matrix (Figure 5e). First, TwINFER indicated that two pairs in the gene co-expression matrix are due to regulation, and the causal relations were determined by the cross-correlations. Both of these interactions have been reported before: *Cebpa* activates *Spi1* [97, 98], and *Spi1* activates *Jun* [99]. Overall, the results indicate that TwINFER can refine the underlying GRN inference.

Next, we found *Gata2* to be highly correlated with five genes (*Egr1, Gfi1, Gata1, Klf1* and *Tal1*), and that the source of these interactions is at least partially due to heterogeneity or a shared confounding factor. The same is inferred for the correlation between *Gfi1* and *Cebpa*; however, here cross-correlations are shown to be insignificant. Lastly, it was previously suggested that *Cebpa* activates *Gfi1* [96], but our analysis suggests that the observed correlations may be of non-regulatory origin. Note that we repeated the check for heterogeneity on day 4, as in bifurcating systems it can emerge later on (Figure E12), and flagged interactions that were inferred as heterogeneous on either of the days. Our predictions could be experimentally tested in future studies.

### Limitations and extensions

TwINFER explicitly relies on the observation that sister cells exhibit greater similarity than random cell pairs drawn from the population [83, 91–93]. Implicitly, we assume that the post-division transcriptome and proteome fully determine the resulting transcriptional steady state (that is, statistically speaking). Therefore, the framework is best suited for analyzing cells undergoing symmetric partitioning, as was done in our simulations in this study. Consistent with this assumption, clonal twins in the LARRY dataset (Figure 5b) and other published barcoding studies [84, 100] exhibit greater transcriptomic similarity than random cell pairs. Nevertheless, the performance of TwINFER will be hampered when partitioning is highly variable, and one should be cautious not to apply this framework to analyze cells exhibiting asymmetric partitioning without the proper extension of the theory herein (see Methods and Figure E13 for demonstration that TwINFER still performs well under symmetric binomial partitioning). Besides, the implicit assumption that transcriptional steady state is determined solely by the initial transcriptome and proteome warrants examination. Higher-order hierarchies of stochastic gene regulation may exist, such as nucleosome positioning [10]. If operating on timescales comparable to TF activity, these could act as confounding hidden variables (see discussion in Ref. [101]), inducing or masking gene-expression correlation.

The application of TwINFER to a dynamic system undergoing differentiation, as was done in this study, presents conceptual challenges. A key assumption of TwINFER is that the GRN structure is static: its connectivity, interaction types (activation/repression), and directionality do not change over time. If hidden variables alter the GRN structure itself during differentiation, our framework will fail. However, if hidden variables represent extrinsic noise affecting gene expression and interaction parameters without changing network structure (e.g., RNA polymerase abundance or the stage in the cell cycle [102]), we have a heterogeneous system of the kind accounted for in Figure 2. In fact, TwINFER will remain useful in such a scenario, additionally allowing us to infer whether regulation occurs on top of heterogeneity. To demonstrate this point, we show the simulation of a sample that gradually bifurcates into two transcriptional states (Figure E12).

## Discussion

The advent of modern synthetic single-cell barcoding [83–88] and other state-of-the-art lineage tracing techniques [103] motivated us to develop TwINFER which leverages mutual information stored in twins. TwINFER addresses two outstanding challenges in correlation-based inference of GRNs: accounting for heterogeneity in cell states and inferring causal relations. We also addressed the longstanding problem of false positives in triplet motifs. Specifically, fan-out and feed-forward loops, commonly observed biological network structures, are frequently misclassified as regulated mutual motifs, but we demonstrated that these motifs are distinguishable with twin information. We conclude this study by performing a proof-of-concept validation of our framework on experimental barcoded single-cell RNA sequencing data.

Modern inference studies often integrate transcriptomics with additional types of information, as surveyed in the introduction [48, 56–62, 64, 65]. We stress that twin information is a distinct and valuable source of data, not merely an analysis pipeline pertaining to the transcriptome. As such, it can be readily integrated into existing multi-omics and multi-modal frameworks. In fact, as twins tend to be similar beyond the transcriptome level, e.g., in proteome and chromatin state, we envision that similar procedures can be applied to extract additional information from both single-cell barcoded multiomic and lineage-resolved imaging datasets, which are beginning to be reported [104, 105].

By compiling experimental data from measurements accumulated over the past two decades [17, 18, 74– 77, 79, 80], we incorporated both elaborate dynamic details—including bursty gene expression and combinatorial non-linear TF-gene interactions—and biologically relevant parameter ranges for the mammalian context into our simulation model. This underscores the power of quantitative measurements to guide theoretical and computational efforts. In turn, our work here informs on areas where gaps remain. For instance, most burst frequency measurements, particularly indirect ones, analyze regulated and unregulated genes together, obscuring the basal bursting rate and the regulatory effect on this rate. Moreover, the commonly used two-state telegraph model to estimate these rates does not account for regulation. Therefore, applying such a model to regulated genes at steady state yields only effective rates. More elaborate models (e.g., Eq. (E3)) may better characterize such systems.

In the same spirit, simulated sample sizes (6,000 twin pairs) in TwINFER paralleled typical scRNA-seq capabilities. Naturally, increasing the number of twin pairs would enhance statistical power, which is increasingly becoming experimentally feasible with rapidly decreasing single-cell profiling costs. Moreover, here we have separated our cells into two samples, measured at two time points. In principle, with substantially larger sample sizes, cells could be separated into multiple samples, each measured at different time points. Such datasets would contain both within-sample twins and cross-sample twins across multiple time points. In systems where twins faithfully approximate each other’s dynamic behavior from the same initial condition, these time-series data could be leveraged to further improve GRN inference through time-series causal inference methods.

The underpinning of the identity of inheritable fate-determining variables and the exact nature of their crosstalk with the GRNs is the subject of active research. As the experimental picture becomes clearer, future follow-ups can extend the modeling and analysis pipeline presented here to better address hidden variables manifested as extrinsic noise.

## Acknowledgments

YG thanks Ian Dardani for initial discussions (2021) during late night science chats in the lab on using twin information for regulatory inference. We thank Goyal lab members, particularly Madeline Melzer, Emmie Grody and Hanxiao Sun for their helpful discussions and feedback. We thank Jaekwon Jung for carefully reviewing the code. We thank Feinberg Information Technology, Northwestern University Information technology, and Quest High Performance Computing Cluster at Northwestern University for providing computing resources. YG acknowledges support from the Pew-Stewart Scholars Program, Burroughs Wellcome Fund Career Awards at the Scientific Interface, Edward Mallinckrodt, Jr. Foundation. We acknowledge support from the National Institute for Theory and Mathematics in Biology (NITMB) through the National Science Foundation (DMS-2235451) and the Simons Foundation (MPTMPS-00005320). YG is a CZ Biohub Investigator. This research benefited from the National Science Foundation through the Center for Living Systems Ignite Award (grant no. 2317138). KA and YS were supported by funding from YG. YZ was supported by a summer grant from the Baker Program in Undergraduate Research, administered by Weinberg College of Arts and Sciences at Northwestern University. BKS acknowledges support from NUCATS T32 and funds to YG. IB was supported by grants to CM and YG.

## Author Contributions

KA, YS and YG conceived and designed the study with input from CM. KA designed and performed the majority of the computational experiments and analyses with input from YZ, YS and YG. YZ performed the computational experiments and calculations for the triplet motifs identification section with input from KA, YS and YG. BKS formulated the linear model and its simulation; YZ extended it and wrote the SM with YS. IB and KA applied TwINFER to the LARRY dataset, with input from YS, YG and CM. YS, KA, YZ and YG prepared the figures and tables. YS and YG wrote the manuscript with input from all authors.

## Data and code availability

This paper analyzes an existing, publicly available dataset [83]. All code for the analyses in this manuscript has been deposited at: https://github.com/GoyalLab/TwINFER. All results were reproduced independently by a non-author (J. Jung) of the paper. We also provide simulation and plot data for all figures in the paper at this Google Drive [link](https://drive.google.com/drive/folders/1apg1QFkGD_QGuxIaUuTUs-l7r6ofpGsA?usp=drive_link).

## Methods

### Modeling regulatory interactions

#### Pairwise regulatory interaction model

Let X and Y be two interacting genes such that gene X encodes for a transcription factor (TF) that regulates gene Y, as illustrated in Figure E1. The model consists of two components: gene expression and regulation.

Gene expression occurs in transcriptional bursts. Specifically, each gene transitions between two states, ‘off’ and ‘on,’ and is transcribed only in the on state. The time spent in a particular state is exponentially distributed with rates *k*_on_ or *k*_off_, respectively. At steady state, the fraction of time spent in the on state is described by the bursting fraction

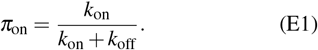

A gene is transcribed into mRNA at a rate 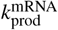, which is in turn translated into a TF at a rate 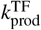. The TFs and mRNAs are degraded at rates 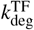 and 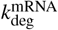, respectively.

Generally, a transcription factor ca regulate the levels of its target gene by modulating the rate of burst initiation (burst frequency) or the rate of burst inactivation of the gene [20]. Here, we model the regulatory effect of a transcription factor as modulation of the burst frequency [11, 24, 27, 106–109]. The type of interaction (activation/repression) determines whether the burst frequency is increased or decreased. For example, if gene X activates gene Y, then it increases the burst frequency of gene Y. The type of interaction is accounted for by the indicator function

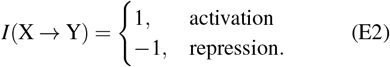

The effective burst frequency of gene Y is given by

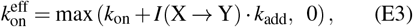

where 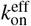is always nonnegative. In the above equation, *k*_add_ is the change in the burst frequency due to regulation. It is a function of the protein levels of gene X, denoted TF_*x*_. The interaction is captured by the Hill function

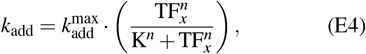

where *n* is the Hill coefficient, K is the Hill constant, and 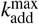 is the maximum possible change in the burst frequency due to an increase in TF_*x*_. More specifically, the steepness of the transition between no regulation and maximal regulation is determined by the Hill coefficient *n*. The Hill constant K is the level of TF_*x*_ at which the change in the burst frequency is half the maximum possible change, and determines the range of levels of TF_*x*_ over which gene Y is sensitive to fluctuations in gene X.

#### Combinatorial regulatory interactions

Let us generalize the above model of pairwise regulatory interaction to a system with multiple interacting genes. In this case, each gene can be regulated by multiple genes simultaneously. We account for the combinatorial regulation with a generalized Hill function following Ref. [81]. For example, in the case of two activators, 1 and 2, the change in the burst frequency of the target in Eq. (E4) can be generalized as

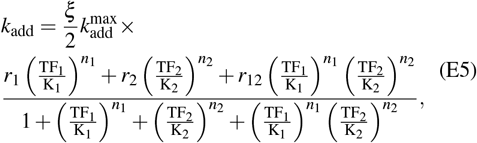

where the way in which the two inputs are integrated to regulate their target is determined by the values of *r*_1_, *r*_2_, and *r*_12_: An ‘OR’ gate is obtained when *r*_1_ = *r*_2_ = *r*_12_ = 1; An ‘AND’ gate is obtained when *r*_1_ = *r*_2_ = 0 and *r*_12_ = 1; and the additive case is obtained when *r*_1_ = *r*_2_ = 1 and *r*_12_ = *r*_1_ +*r*_2_ = 2. If we require that one TF has a stronger interaction than the other, we can set different *r*_1_ and *r*_2_. The prefactor *ξ* is set to 1, 4*/*3, and 4 for additive, OR and AND, respectively. In this way, the definition of K can be maintained such that at this concentration 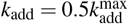. Note that at saturation (where TF concentrations are effectively infinite), for the additive combinatorial interaction we get 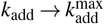, as was the case with the pairwise interaction. For the OR and AND gates we obtain 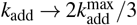 and 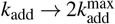, respectively.

The additive interaction also admits an intuitive form

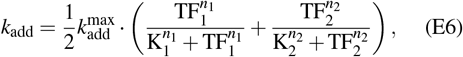

where two pairwise interactions of the form in Eq. (E4) are summed (assuming equal weights). This result is easily generalizable to account for any number of regulators.

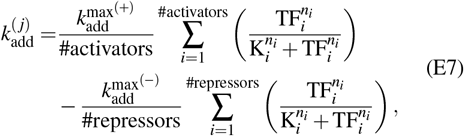

where 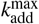 takes different values for positive (+) or negative (−) interactions.

### Model parameters

We require six parameters to model the expression of a gene and three parameters to model a regulatory interaction.

Let us start with the gene expression parameters. In several previous studies of mammalian cells, these parameters were measured directly or inferred from experimental data [17, 18, 74–80]. To determine biologically relevant parameter ranges for our simulation, we compile the values across the different datasets and use the fifth to ninety-fifth percentiles. In the GitHub repository of this paper, we provide the code and the datasets used to define the ranges. Table E1 summarizes the ranges and medians of the parameters, along with references to the corresponding datasets used to obtain them. Similar ranges have been used for modeling bursty gene expression in other studies [71, 110]. Note that we calculate the mRNA and protein degradation rates 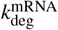 and 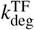 from half-life measurements, assuming first-order decay.

In Table E1, it can be seen that the estimated values of *k*_on_ and *k*_off_ span several orders of magnitude. However, the estimated bursting fraction *π*_on_, defined in Eq. (E1), is constrained to a subspace of all possible combinations of *k*_on_ and *k*_off_. Indeed, the bursting fraction is often very small, reflecting the bursty nature of gene expression. Therefore, a pair of *k*_on_ and *k*_off_ must be constrained to the subspace that ensures bursty expression. In practice, when performing a parameter scan, we first sample *π*_on_ and *k*_on_, and then compute *k*_off_ using Eq. (E1). In this manner, we guarantee that all sampled parameters are within their range.

Let us now consider the regulatory interaction parameters, summarized in Table E2. The motivation behind the ranges used for these parameters was two-fold: To capture diverse regulatory conditions, and to ensure gene Y’s sensitivity to fluctuations in TF_*x*_ across the population.

We used a range of values of *n* to sample different steepness levels of the Hill function representing diverse regulatory interactions, consistent with prior works [23, 110].

The parameter 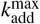 is not estimated in the experimental datasets. This is because the two-state telegraph model, which does not include regulation, is routinely used for their estimation (which also means that the reported *k*_on_ values must be taken with a grain of salt). Nonetheless, it has been reported for some model genes that the impact of regulation results in a four-to ten-fold increase in *k*_on_ [25, 27, 112].

We set the Hill constant K to the mean TF_*x*_ level under steady-state conditions so that the TF_*x*_ levels lie in the steep region of the Hill function

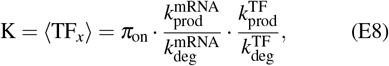

where all the parameters belong to the regulating gene X. Note that at steady state, by the definition of K, Eq. (E4) yields *k*_add_ = 0.5 · 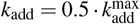.

For a regulated gene Y, the bursting fraction *π*_on_ is a function of 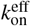, which in turn depends on *k*_add_, as described in Eq. (E3). Finally, Eq. (E4) tells us that *k*_add_ is a function of TF_*x*_. To make that dependency explicit, we write

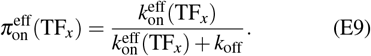

If we are interested in ⟨TF_*y*_ ⟩, Eq. (E8) must be modified accordingly

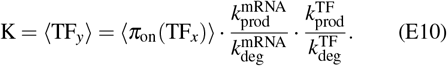

Nota bene, *π*_on_ is a concave function of a random variable. Thus, from Jensen’s inequality [114],

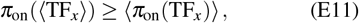

and one cannot simply exchange between the two. This introduces a complication, as Eq. (E10) must be evaluated numerically by simulating the system, unlike Eq. (E8) which can be computed analytically. Thus, if we want to set K as the steady state of a regulator that is itself being regulated, we have to simulate the network up to that interaction to find the steady state concentration of the regulator numerically. When dealing with a gene cascade, for example, we must perform a hierarchical series of simulations, each time adding another downstream edge. That said, in our hands, the error introduced by using Eq. (E10) to evaluate the steady state in this case is only a few percents and thus inconsequential (see Figure E6c).

To date, the parameters of the pairwise regulatory interaction model in Table E2, or any other interaction model for that matter, have not been directly measured or inferred. Therefore, we had to reason which values would constitute a typical interaction strength. On the one hand, the interaction has to be strong enough to result in correlations that persist beyond the noisy gene expression. On the other hand, the regulation must not be too strong such that the regulated gene ceases to be bursty altogether, i.e., reaches a saturated regime where *π*_on_( ⟫TF_*x*_ ⟩) = 1. In fact, the correlations are expected to drop in the saturated regime, as higher concentrations of the TF will yield increasingly diminishing effects. In Figure E6a-b we scan the relative sizes of 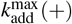 and 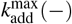 compared to the median *k*_on_. All other parameters were also set to the median values in Table E1. Indeed, it can be seen that the magnitude of the gene-expression correlation *ρ* gradually increases until it saturates and the trend reverses. We chose the default value for our simulation so that the interactions are strong, but still well below that of the saturated regime. Likewise, in Figure E6c we show that our choice for K = ⟨TF ⟩ falls well within the highly correlated, yet unsaturated, regime.

**TABLE E1.** Bursty gene expression model parameters. Medians and ranges were extracted from published studies with direct measurements, as well as inference schemes relying on indirect measurements, of various mammalian cells. The ranges are determined by the fifth to ninety-fifth percentiles of the compiled data for each parameter.

| Parameter | Description | Range | Median | Units | References |
| --- | --- | --- | --- | --- | --- |
| $k_{\text{on}}$ | Rate of burst initiation | 0.01 – 3 | 0.66 | $h^{-1}$ | [17, 18, 74] |
| $k_{\text{off}}$ | Rate of burst inactivation | 0.06 – 55 | 8.6 | $h^{-1}$ | [17, 18] |
| $\pi_{\text{on}}$ | Bursting fraction | 0.002 – 0.4 | 0.07 | – | [17, 18] |
| $k_{\text{prod}}^{\text{mRNA}}$ | Transcription rate | 0.2 – 60 | 2 | $\text{mRNA} \cdot h^{-1}$ | [17, 18, 75, 79] |
| $k_{\text{prod}}^{\text{TF}}$ | Translation rate | 19 – 2700 | 560 | $\text{protein} \cdot \text{mRNA}^{-1} h^{-1}$ | [75, 79] |
| $\tau_{1/2}^{\text{mRNA}}$ | mRNA half life | 0.6 – 17 | 4 | $h$ | [18, 76–79] |
| $\tau_{1/2}^{\text{TF}}$ | Protein half life | 7 – 200 | 45 | $h$ | [79, 80] |

**TABLE E2.** Pairwise regulatory interaction model parameters. Each regulatory interaction requires three additional parameters, where 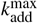 takes different values for positive (+) and negative (−) interactions. The range for possible 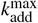 values is a function of *k*_on_ (linear scaling).

| Parameter | Description | Range or Value | Simulation Default |
| --- | --- | --- | --- |
| $k_{\text{add}}^{\text{max}}(+)$ | Activation strength | $(0.5 - 10) \cdot k_{\text{on}}$ | 6 |
| $k_{\text{add}}^{\text{max}}(-)$ | Repression strength | $(0.5 - 2) \cdot k_{\text{on}}$ | 0.8 |
| $n$ | Hill coefficient | 0.5 – 5 | 2 |
| $K$ | Hill constant | Steady-state of TF | – |

### Simulation algorithm

In the GitHub repository of this paper, we provide the code for our simulation and analysis pipelines.

Based on the gene expression and interaction model described above, the simulation provides the gene expression profiles for an arbitrary GRN. Specifically, the simulation output is the mRNA and protein levels of the *N* twin pairs, measured at time points specified by the user.

The simulation input is the number *N* of twin pairs to be simulated, the gene expression parameters, the regulatory interaction matrix (connectivity matrix), and the regulatory interaction parameters for each interaction.

The simulation tracks the cell state of each cell, comprising the bursting state, mRNA levels, and protein levels of all genes in the regulatory network. Cell states evolve through three processes: stochastic switching of bursting states, the production of mRNA and proteins, and the degradation of mRNA and proteins. Each of these processes has a reaction propensity, which determines the probability of the corresponding event occurring during an infinitesimal time interval d*t*. As previously used in Ref. [23] and Ref. [110], we employ Gillespie’s next reaction method [113] to simulate the evolution of cell states based on the state-dependent reaction propensities.

The simulation starts with *N* mother cells, each with zero mRNA and protein counts and with all genes initially inactive. The mother cells are allowed to evolve until the population-level cell states reach a steady state (it takes a thousand hours or more with our parameters). We test for steady state by comparing the mean transcript level with the analytical expressions in Eqs. (E8) and (E10). After steady state is established, we generate *N* pairs of identical twin cells, one pair for each mother cell. The daughter cells are separated into the two samples, assuming well mixing. For example, when separating 6, 000 twin pairs, we end up with 1, 500 within-sample twin pairs in each sample, and with 3, 000 cross-sample twins pairs. We then start the clock. The cells evolve their cell state independently until their measurement time (*t*_1_ for Sample 1 and *t*_2_ for Sample 2), where we record the transcript and protein counts.

All simulations presented in the main text were performed with perfect partitioning of the transcriptome and proteome upon division of mother cells. In Figure E13, we show that TwINFER performs well even when binomial partitioning [115] is simulated. Note that we simulated symmetric binomial partitioning, i.e., the probability that a transcript or a protein molecule ends up in either daughter cell is the same, *p* = 0.5. The variance is given by *p*(1 − *p*)*c*, where *c* is the number of copies of the partitioned molecule in the dividing cell. By varying *p* in our code, one can explore asymmetric binomial partitioning.

### Statistical tests employed in Figure 2

Figure 2a presents a flowchart with three stages, each involving a different statistical test. We will further elaborate on each test.

Test for stage I: We perform a permutation test, where we compute 10, 000 scrambled gene-expression profiles (Figure E3). The null hypothesis is that the observed correlation *ρ* is drawn from a distribution obtained from the many scrambles. We reject it with a correlation threshold corresponding to the desired p-value.

Test for stage II: We compute *ρ*_Δ_ and 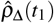. While for each cell there is only a single possible twin counterpart, random pairs can be drawn in many different ways. We thus perform another permutation test, drawing 10,000 random pairings (Figure E4). This results in a distribution for *ρ*_Δ_. We then perform a z-test to determine whether 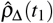 is drawn from this distribution. We conclude cell-state heterogeneity if the z-score is below the desired threshold.

Test for Stage III: Unlike the previous two stages we require multiple repetitions of the experiment or sub-sampling of the data in order to ensure statistical significance. Thus, we perform a Wilcoxon signed-rank test between 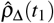 and 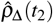. This is a paired comparison test. A significant increase corresponds to Scenario 3, otherwise we infer Scenario 4.

### Special analysis considerations

Stage I of the flowchart in Figure 2 involves the calculation of *ρ*, the ordinary gene-expression correlation across a population of single cells. Since *ρ* is time-independent, we combine data from both samples to gain more statistical power. Note the underlying assumption that the GRN itself does not change during the time interval [*t*_1_, *t*_2_]. If it does, a more careful analysis is needed.

The same reasoning is applied in Stage II, where we compute the time-independent *ρ*_Δ_ across random pairs of cells, again merging the data from both samples.

In Figures 2, E3 and E5, the interaction type is known to be either activation or repression, and determines the sign of the gene/cross correlations (positive and negative, respectively). The distributions of correlations from shuffled gene-expression profiles are symmetric around zero (in fact, these are normal distributions with mean zero). We thus computed one-tailed (directional) p-values. For example, a 0.01 p-value corresponds to the 1st percentile for repression and the 99th percentile for activation. In Figures 3 and 5, we do not assume to know the type of interactions. Therefore, we computed two-tailed (non-directional) p-values. To do so, we computed the one-tailed p-values for both sides, *p*^−^ and *p*^+^, and then used *p* = 2 · min(*p*^+^, *p*^−^).

### Scanning the parameter space to define inference capabilities and limitations

The results presented in Figures 2 and 3 and discussed in the main text are from in silico experiments performed with the median values in Table E1 and the default values in Table E2. These values are not necessarily the ideal values for inference, yet TwINFER had a perfect F1 score of 1. To further challenge TwINFER and better define its inference capabilities and limitations, we scanned the parameter space restricted by the ranges in Tables E1 and E2. The results are plotted in Figure 2i-k.

Recall the flowchart shown in Figure 2a, summarizing the tests performed to determine whether a correlation emanates from a regulatory interaction or from the existence of multiple states. In the main text, whenever we simulated Scenarios 3 and 4, the mother cells were assigned into two equally sized groups, each with a different bursting frequency *k*_on_ shared by all genes in that state, 0.12 *h*^−1^ and 1.66 *h*^−1^. This is an intuitive way to obtain heterogeneous transcriptional states while keeping all the rest of the parameters the same across the cell population. The very different bursting frequencies resulted in unmistakably different protein count distributions, with no overlap at all.

However, it is clear that during the parameter scan, the randomly drawn parameter sets for the two states of a gene can happen to be similar enough to result effectively in a single-state, and one should not classify such cases as heterogeneous systems. To prevent such misclassifications, we set a more rigorous criterion for the existence of multiple states: for each gene, we perform the Brunner–Munzel test for the protein count distributions of cells coming from each of the states. Only if *both* genes pass the test (*p <* 0.01), we refer to the system as heterogeneous. When simulating 25,000 parameter sets for the multiple-state scenarios, we discard systems that fail this criterion and continue sampling until 25,000 heterogeneous sets are reached. In Figure E7 we plot the protein count distributions for a representative parameter set that satisfies the criterion and for one that does not.

### GRNBoost2

We used the grnboost2 function available with the Arboreto package (version 0.1.6) to analyze our in silico data (Figures 3 and 4). The details of the package and its installation are available in the documentation on their [website](https://arboreto.readthedocs.io/en/latest/index.html).

GRNBoost2 produces a ranked list of candidate interactions, scored by a feature-importance measure derived from gradient-boosted decision-tree models (inspired by GENIE3) [51, 52]. However, GRNBoost2 does not inform the user on how to threshold the ranked list. In the absence of an optimized protocol for determination of the threshold, different users will choose different thresholds, and thus infer different GRNs.

In this study, we developed our own method for thresholding, based on a Gaussian Mixture Model (Figure E8). We fit a two-component Gaussian mixture model to the distribution of GRNBoost2 importance predictions, separating the interactions into two Gaussian components corresponding to low and high importance. We set the importance threshold at the intersection of the two weighted Gaussian probability density functions, where an edge is equally likely to belong to either component. We acknowledge that, at least in principle, one could devise a better procedure for threshold optimization. We therefore decided to try all possible thresholds and pick the one that performs the best (in terms of the highest F1 score). We refer to this threshold as the upper bound of GRNBoost2. Of course, in practice, one does not know the ground-truth network under question, and so the upper bound is unknown.

### Applying TwINFER to a lineage- and time-resolved scRNA-seq dataset of hematopoietic differentiation

To evaluate our pipeline on experimental data, we used the LSK LARRY dataset, which contains scRNA-seq data from LSK mouse hematopoietic progenitors (Ref. [83]). Cells were collected at 2, 4 and 6 days after lentiviral barcode labeling to capture successive stages of broad myeloid differentiation. Data preprocessing included removing cells with *>* 20% mitochondrial gene content and merging lineage barcodes within a Hamming distance of 3, as reported. Doublets were removed using the singletCode package (Ref. [85]), following the procedure applied to the LARRY dataset in that study. Subsequently, cells expressing fewer than 200 genes were discarded, and gene expression values were normalized to 10,000 counts per cell and log-transformed.

For the analysis, we selected cells from days 2, 4, and 6 to define time points *t*_1_, *t*_2_, and *t*_3_, respectively. The dataset included cell-type annotations for each cell, either as undifferentiated or as one of nine mature cell types. We annotated the fate of day 2 cells by the cell type most frequently observed in day 4 and/or day 6 cells from the clone. If the most frequent cell type at day 4 was undifferentiated, we used the most frequent cell type at day 6, if available. Our objective was to test a GRN implicated in the myeloid differentiation model, as described in Ref. [95]. Therefore, we chose cells that were undifferentiated at day 2, and were not identified to have a dominant lymphoid fate. In other words, we excluded barcodes with a majority of clones categorized as lymphoid or pDC at days 4 and 6. Additionally, we also excluded cells at day 4 and day 6 that were annotated as lymphoid or pDC cell types.

For differential gene expression, we calculated the difference in expression between cells annotated as having a particular fate and all other day 2 cells, following the procedure described in Ref. [83].

All possible cell pairs within each clone were considered, as each clonal pair represents a potential differentiation trajectory. While we acknowledge that these clonal pairs do not necessarily correspond to true twin cells, we included all clonal pairs as twin cells were not identifiable in the dataset. We validated the use of clonal pairs as a proxy for twin pairs by comparing the within-sample PCA distance of clonal pairs and random pairs. As seen in Figure 5b, we found that within-sample clonal pairs were significantly more similar to each other. Therefore, we included clonal pairs between all possible cells found in a particular clone. We had 1547 within-sample clonal pairs for day 2 and 7471 for day 4. We had 2879 cross-sample clonal pairs between these days.

We employed two additional filters. First, we addressed the dynamic nature of differentiation, as the GRN changes over time. We identified a static subnetwork of genes with stable correlations (Figure 5e), i.e., correlations that remained statistically significant and of the same sign throughout the experiment. Second, we observed that a small, lowly expressed sub-group of genes has been measured sparsely and is absent from most measured cells. When performing the shuffle tests for interactions involving these genes, we could fail to obtain the expected normal distribution (under the central limit theorem). To filter such interactions, we plotted Q-Q plots, comparing each distribution of correlations from shuffled profiles with a normal distribution. Fitting the Q-Q plots with the identity line, we excluded interactions with an R-squared less than 0.9. We found that this threshold was enough to eliminate abnormal-looking distributions (three in our case). We label the gene pairs filtered out by either filter as no-regulation.

## Extended Figures

**FIG. E1.**
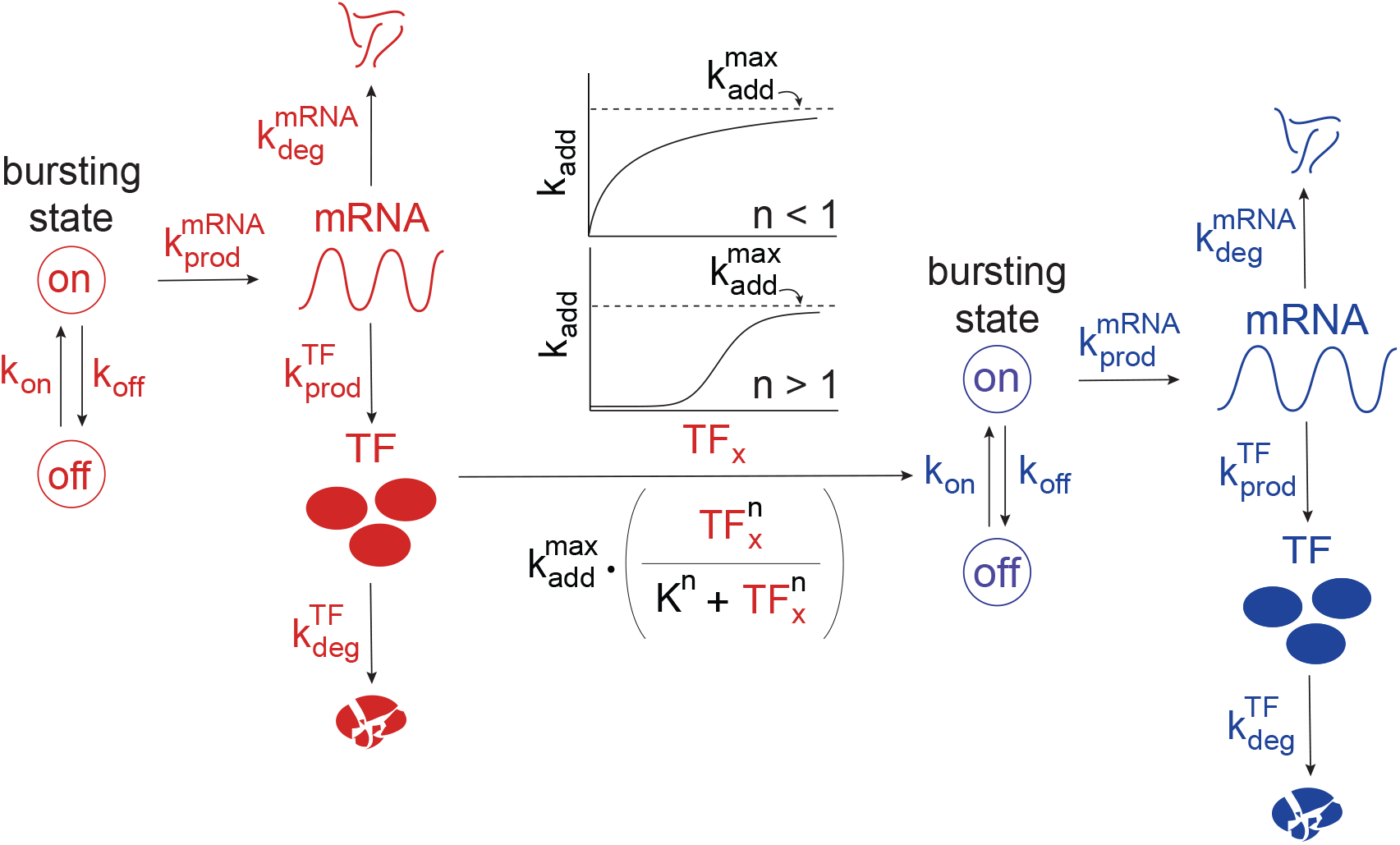
A model of a pairwise regulatory interaction. Gene X (red) regulates gene Y (blue). Each gene transitions between two states, ‘on’ and ‘off’, with corresponding rates *k*_on_ and *k*_off_. Transcription occurs only in the on state, at a rate 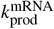. It is followed by protein synthesis at a rate 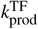. Gene Y is regulated by the modulation of its rate of burst initiation *k*_on_. The strength of regulation increases with the levels of the TF_*x*_, and is modeled as a Hill function with a maximum possible change in rate 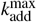, a Hill constant K, and a Hill coefficient *n*.

**FIG. E2.**
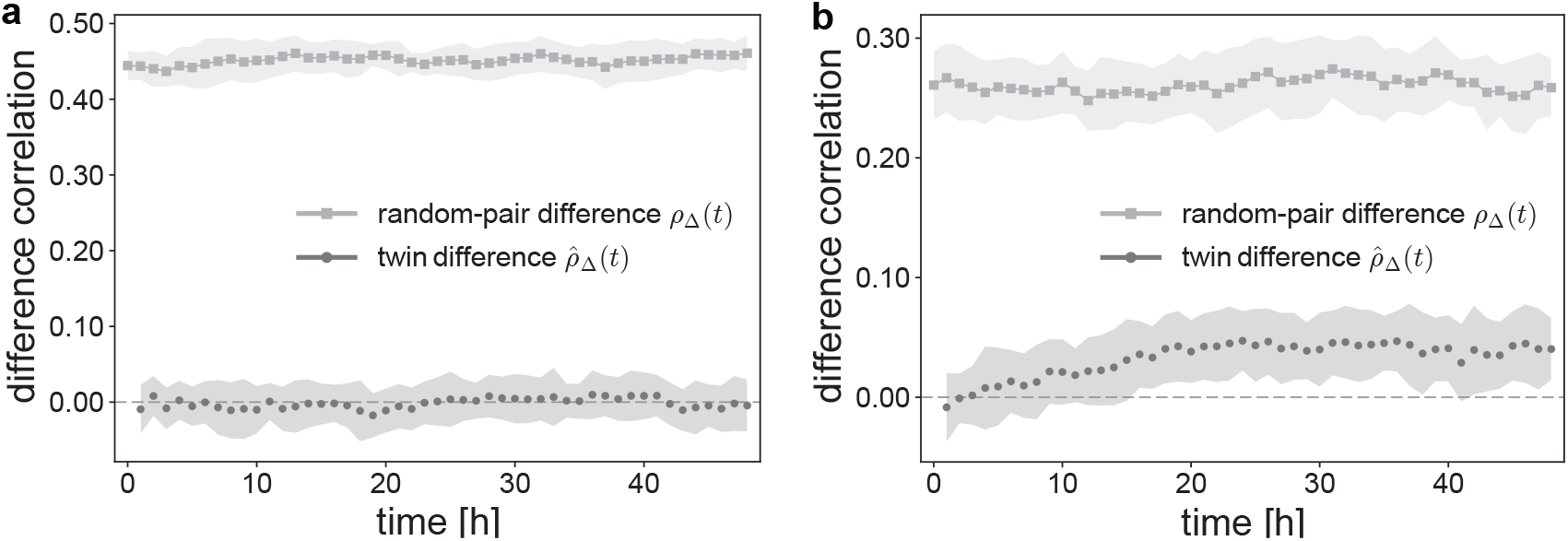
A system of two genes, X and Y, each with two states. Random-pair correlation and within-sample twin correlation of the differences Δ*x* and Δ*y*, denoted *ρ*_Δ_ and 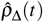 respectively, plotted here as a function of time *t*. **(a)** X and Y do not interact, namely there is no regulation. **(b)** Gene X activates gene Y. Before division, the mother cells were assigned to two equally sized groups, each with a different value of *k*_on_, 0.12 *h*^−1^ and 1.66 *h*^−1^ (the Brunner–Munzel test for protein count distributions in the two different cell states yielded *p* = 0). All other parameters were set to the median and default values in Tables E1 and E2, respectively. The standard deviation bands were computed from 20 repetitions, each with 6, 000 twin pairs.

**FIG. E3.**
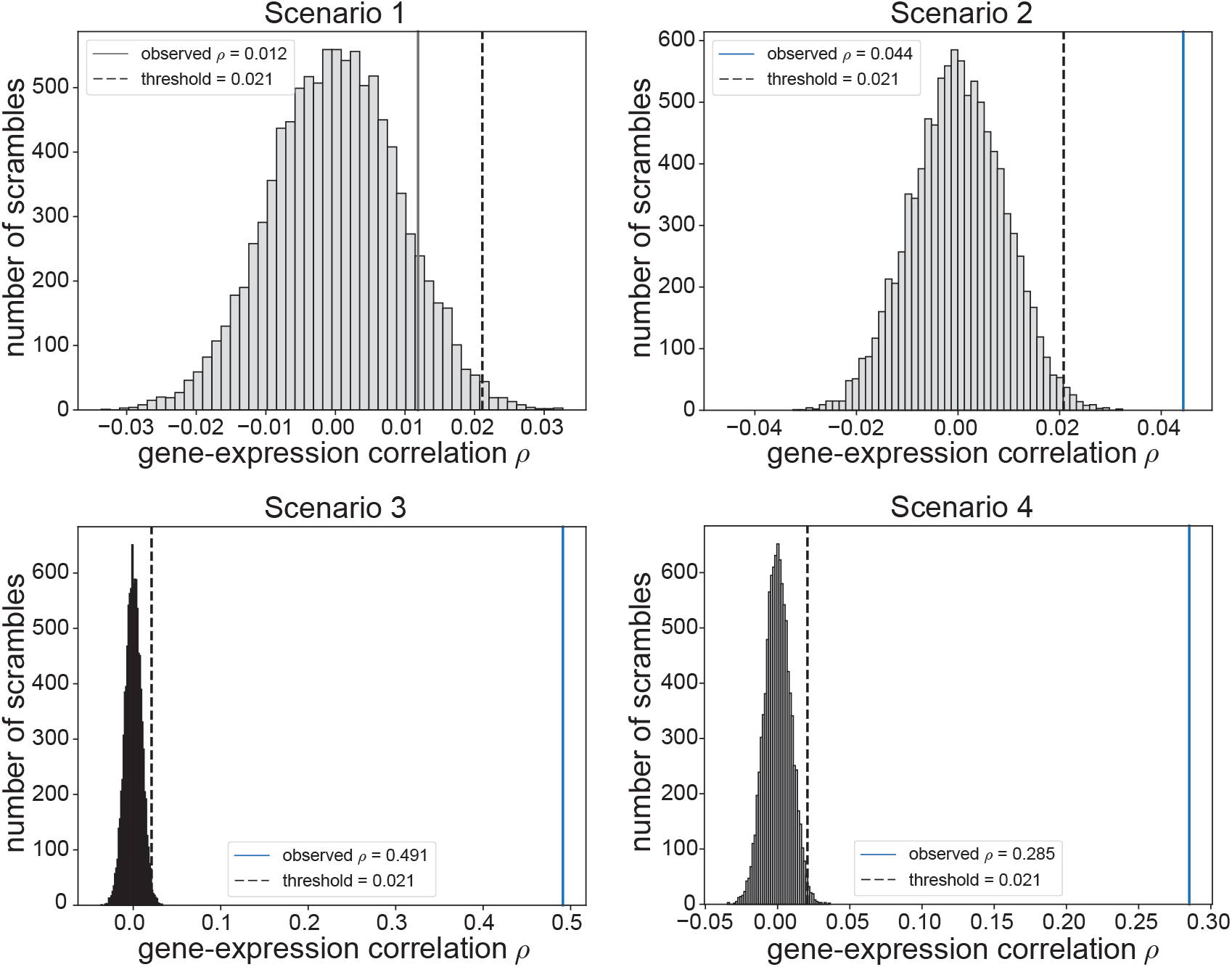
Statistical test for stage I of the flowchart in Figure 2a. Distributions of the gene-expression correlations *ρ* from 10, 000 random scrambles of the original gene-expression profile, for all four scenarios. Each subplot is a representative distribution produced for one out of the twenty repetitions performed in Figure 2d. The threshold correlations, indicated by dashed vertical black lines, correspond to a one-tailed *p* = 0.01. The actual (not scrambled) correlations *ρ* are indicated by the vertical lines and are colored blue if they pass the threshold.

**FIG. E4.**
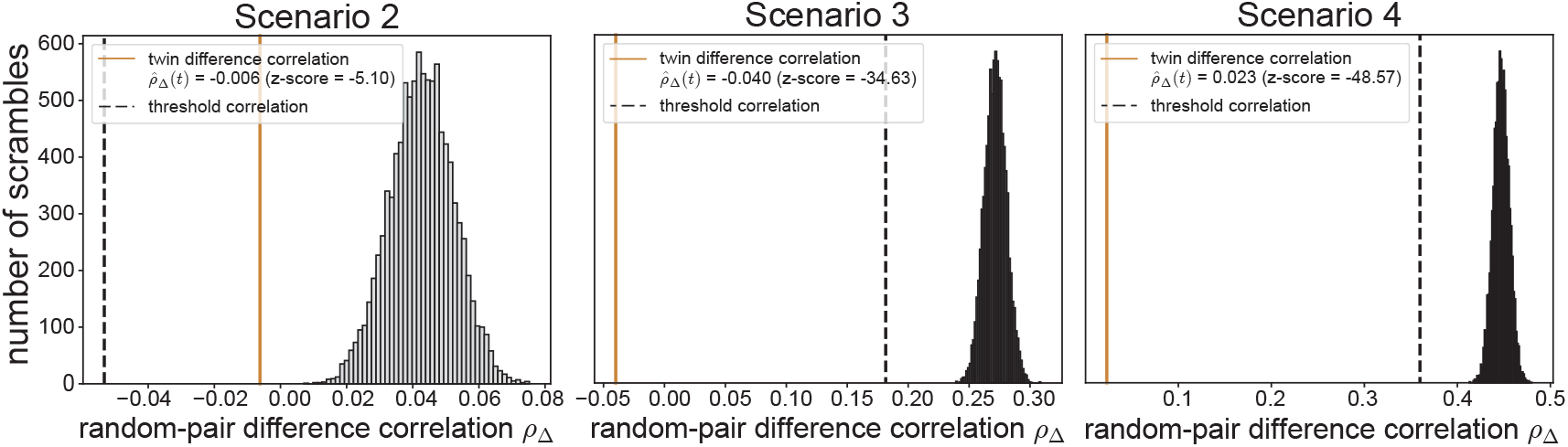
Statistical test for stage II of the flowchart in Figure 2a. Distributions of random-pair difference correlations *ρ*_Δ_ from 10, 000 different possible random pairings of the gene-expression profile, for the three possible scenarios (Scenario I is eliminated in Stage I). Each subplot is a representative distribution produced for one out of the twenty repetitions performed in Figure 2f. The threshold z-scores, indicated by dashed vertical black lines, correspond to a z-score of −10. The twin difference correlations 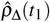 are indicated by the vertical orange lines.

**FIG. E5.**
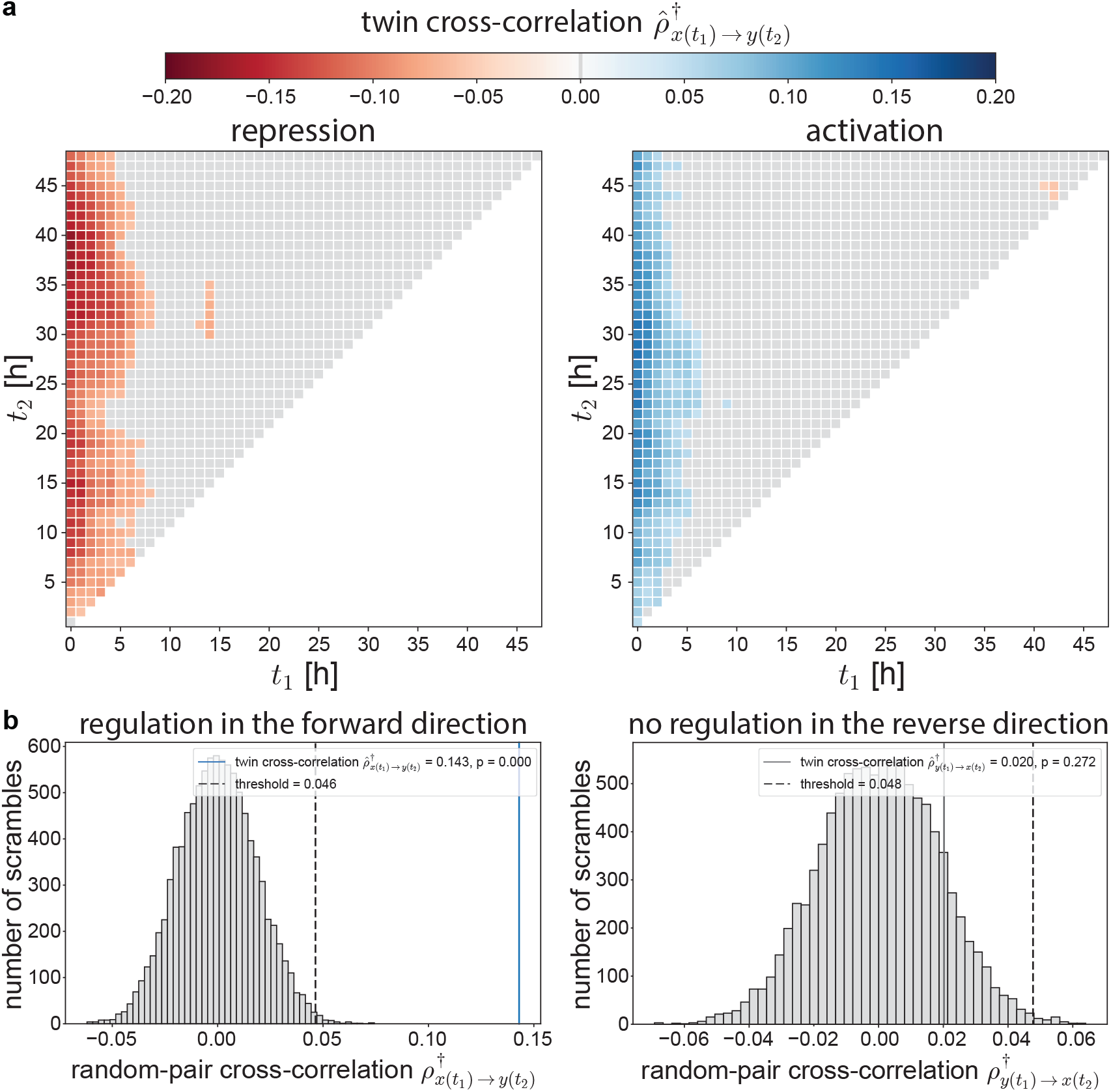
**(a)** Robustness of TwINFER under different measurement times. In the main text, measurements were taken at *t*_1_ = 1 and *t*_2_ = 20 hours. Here we keep the simulation parameters the same (i.e., the median and default values in Tables E1 and E2), and vary *t*_1_ and *t*_2_. We compute the cross-correlations for unidirectional activation and repression. Gray squares represent insignificant interactions—working with these pairs of *t*_1_ and *t*_2_ will result in false negatives. One can observe that the window for *t*_1_ is limited, it is best to measure Sample 1 within a few hours after the division of the mother cells. In contrast, the window for *t*_2_ is much wider and spans tens of hours, with best performance when measuring Sample 2 around ∼ 15 − 35 hours after the division. **(b)** Statistical test for the significance of a directional interaction X → Y. We plot distributions of random-pair cross-correlations 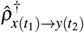 (left panel) and 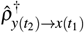 (right panel) from 10, 000 different possible random pairings of the gene-expression profile for the representative simulation of X → Y activation shown in Figure 3. The left panel shows regulation in the forward direction, corresponding to 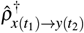, and indeed the twin cross-correlation (blue vertical line) is significantly different than the scrambles (*p* = 0). Conversely, the right panel demonstrates that since there is no regulation in the reverse direction, corresponding to 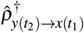, the twin cross-correlation (gray vertical line) is *not* significantly different than the random-pair scrambles correlations. Note that the statistical tests in this figure are one-tailed, as we know the type of the interactions (activating/repressing). When this information is not available, as is the case in Figures 3 and 5, we perform a two-tailed test.

**FIG. E6.**
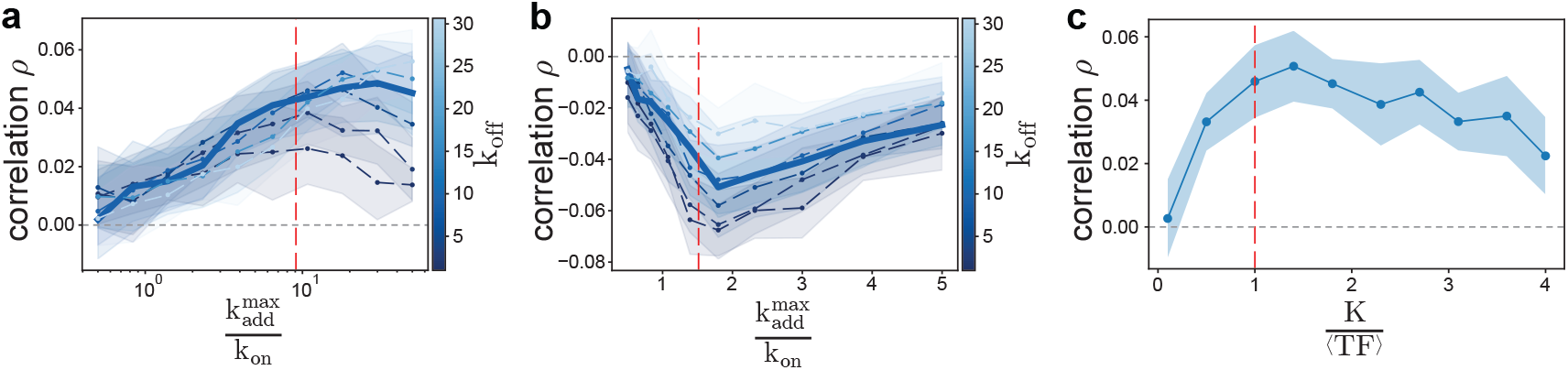
Choice of the interaction model parameters. We plot the gene-expression correlations *ρ* vs. (**a-b**) the ratio between the maximal possible change in burst frequency 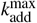 and the median value for *k*_on_; and (**c**) the ratio between the Hill constant K and the steady state transcription factor concentration ⟨TF ⟩. The vertical red dashed lines correspond to the default ratios set in the simulations. Subplots (**a**) and (**b**) are for activation and repression, respectively. Each curve in (**a**) and (**b**) represents a different value of *k*_off_, where the bold curve is the median. All other parameters were set to be the median values in Table E1. The standard deviation bands were computed from 20 repetitions, each with 6, 000 twin pairs.

**FIG. E7.**
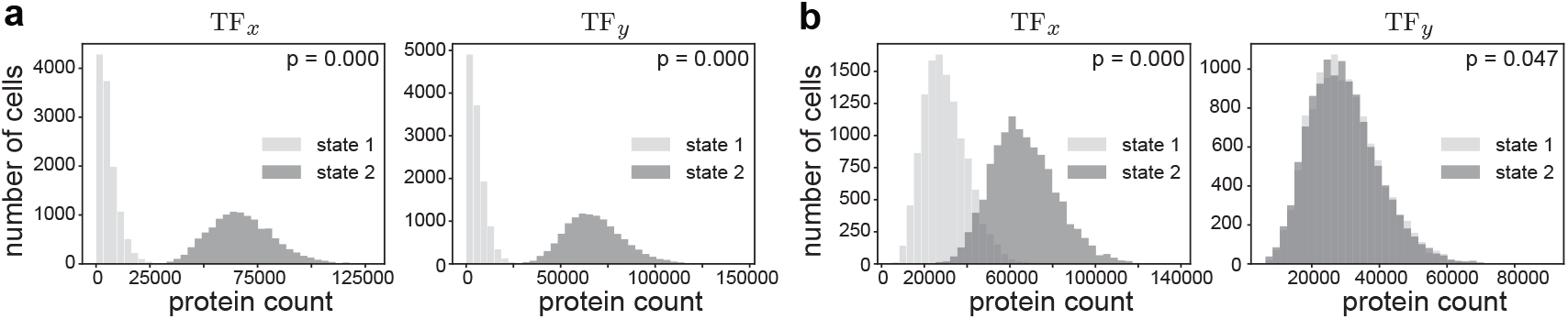
As part of the parameter space scan, we simulated 25,000 different heterogeneous systems, where both the genes X and Y have two transcriptional states. To create two states, the mother cells were assigned into two equally sized groups, each with a different parameter set for each gene in that state. The randomly drawn parameter sets can happen to result in protein count distributions that are statistically indiscernible. Such cases are effectively a single state and should not be classified as a heterogeneous system. We thus perform the Brunner–Munzel test for the protein count distributions of each gene’s two states. Only if both genes have significantly different distributions (*p <* 0.01) do we deem the system heterogeneous. We kept simulating systems with different parameter sets until we reached 25,000 heterogeneous systems that fulfilled this criterion. In **(a)** we plot the protein count distribution for both genes of the representative heterogeneous system (*p* = 0) from Figure 2, i.e., with all genes in a state sharing a bursting frequency *k*_on_ of either 0.12 *h*^−1^ or 1.66 *h*^−1^, and where all other parameters are the median and default values from Tables E1 and E2, respectively. In **(b)** we plot a simple representative system with an effective single state (*p* = 0.047 for gene Y). In this example, all parameters for all genes are the median except for one state of gene X which has a bursting frequency 1.66 *h*^−1^. Evidently, this is not enough to qualify as a heterogeneous system.

**FIG. E8.**
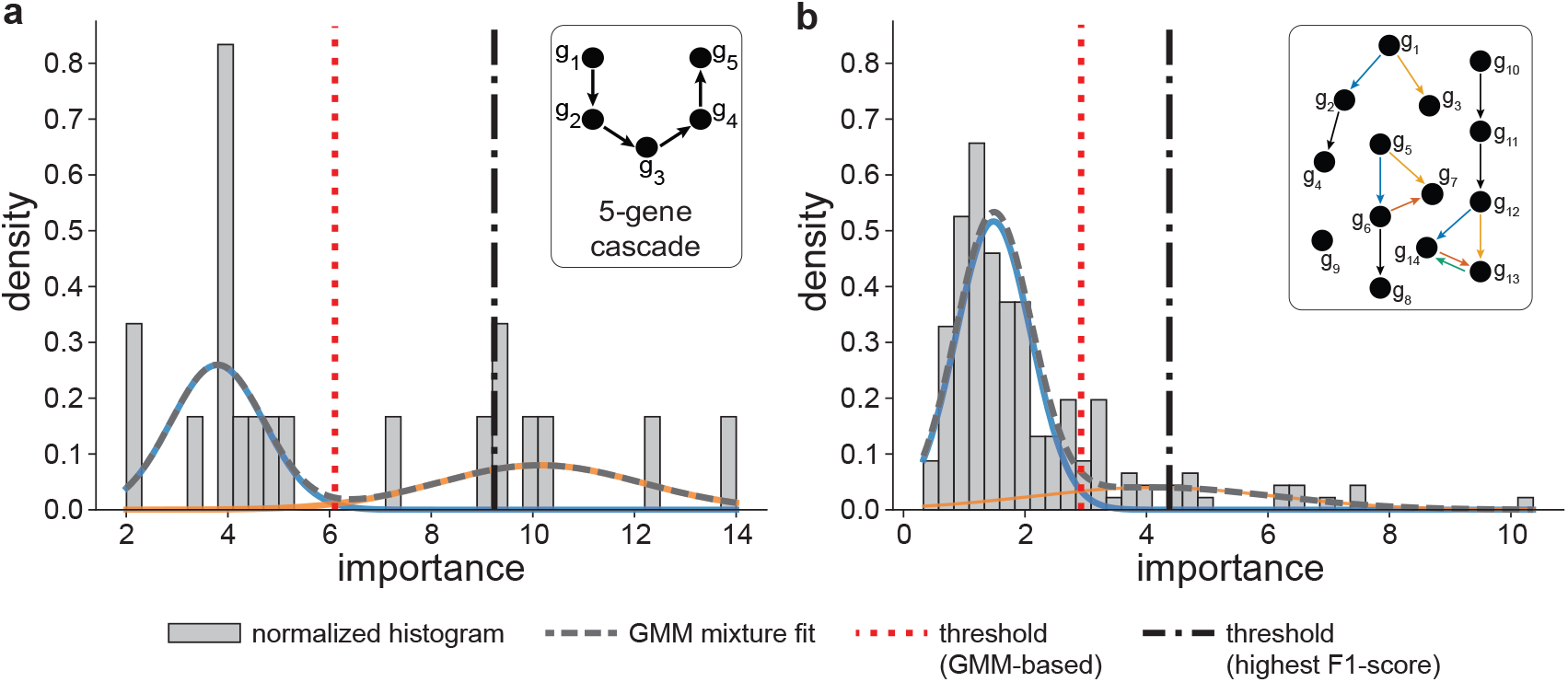
Thresholding the GRNBoost2 ranked list using a Gaussian Mixture Model (GMM). We fit a two-component Gaussian mixture model to the distribution of GRNBoost2 importance predictions to separate the interactions into two Gaussian components corresponding to low and high importance (blue and orange curves, respectively). We set the importance threshold at the intersection of the two weighted Gaussian probability density functions, where an edge is equally likely to belong to either component. We acknowledge that, at least in principle, one could devise a better procedure for threshold optimization. We therefore decided to try all possible thresholds and pick the one that results in the highest F1 score (dashed dotted line). We refer to this threshold as the upper bound of GRNBoost2. Of course, in practice, one does not know the ground-truth network under question, and so the upper bound is unknown. In **(a)** and **(b)** we analyze representative simulations of the 5-gene cascade from Figure 3 and the 14-gene network from Figure 4, respectively.

**FIG. E9.**
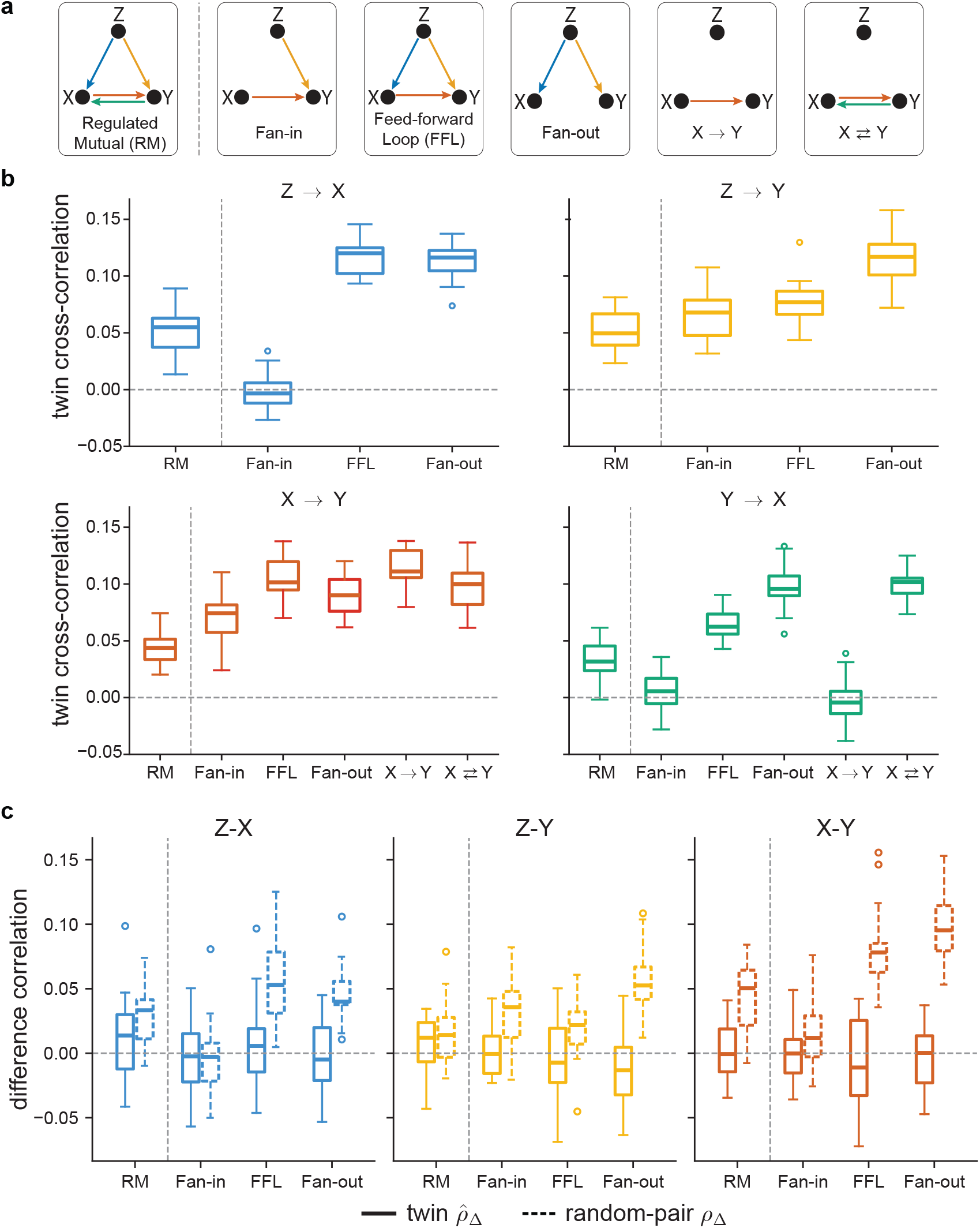
Attenuation of correlations in regulated mutual (RM) are caused by the two embedded fan-in sub-motifs. **(a)** The regulated mutual motif and all sub-motifs embedded within it. **(b)** Twin cross-correlations of RM and its sub-motifs. Out of all sub-motifs, fan-in results in reduced correlations, comparable to the reduction seen in RM. **(c)** The difference correlation between X and Y is completely attenuated in fan-in, but not in FFL and Fan-out, suggesting that fan-in is the cause for the attenuation of the difference correlations in RM.

**FIG. E10.**
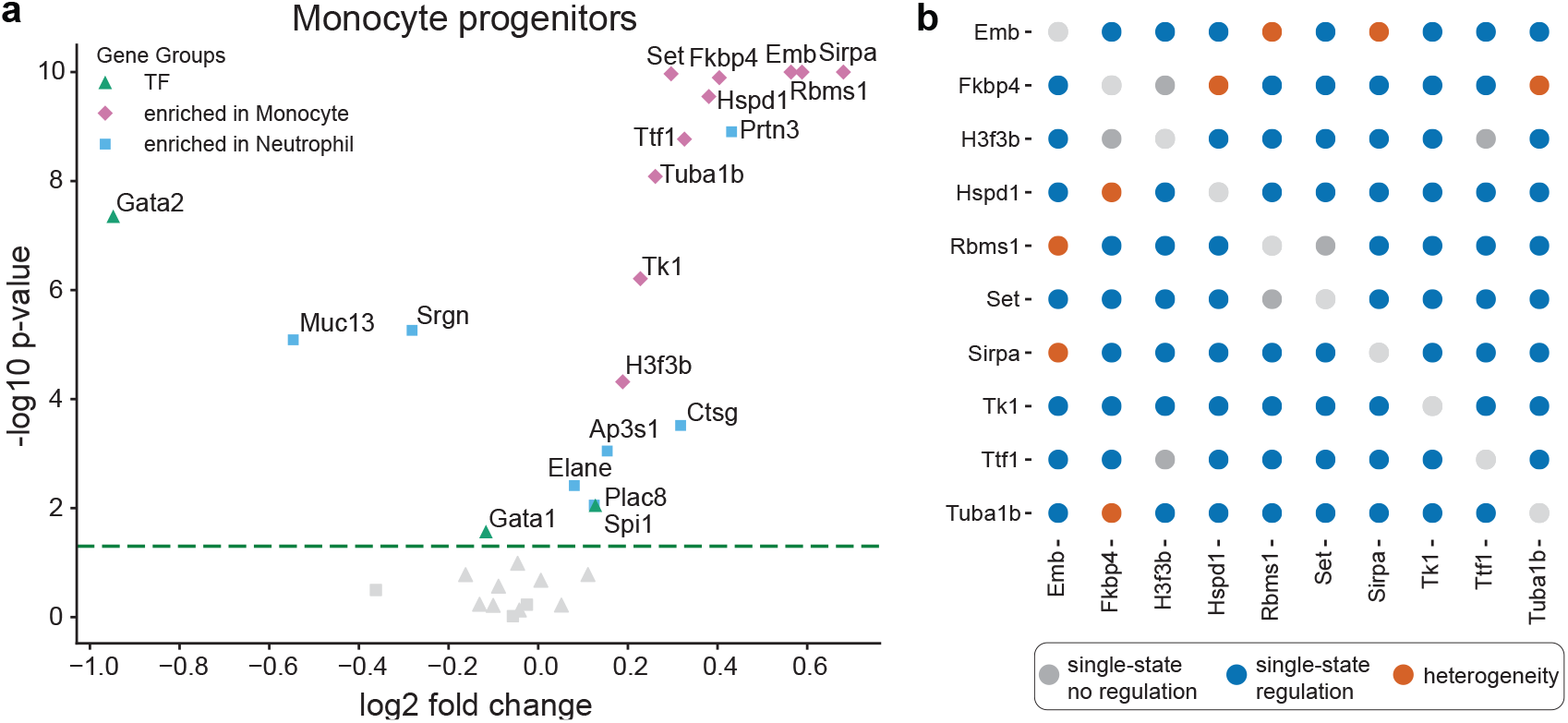
Here we repeat the analysis that was performed in Figure 5c-d but for early monocyte progenitors. **(a)** A volcano plot, corresponding to the analysis in Figure 2G of Ref. [83], but repeated with our protocols. Undifferentiated early progenitors were categorized based on their twin’s fate. The plot identifies genes that are enriched in monocyte progenitors compared to the general population at day 2. We tested three groups of genes. First, a group that was reported to be differentially expressed in monocytes, and another in neutrophils [83] (diamonds and rectangles, respectively). Second, a group of TFs, known to be driving murine myeloid differentiation [95] (triangles). Most TFs were not differentially expressed (gray). In our reproduction, all genes in the monocyte group were found to be significantly enriched in monocyte progenitors. **(b)** For the group of genes enriched in monocyte progenitors, we plot a dot matrix. Each pairing of genes is denoted by a dot, its color representing a different scenario inferred by following the flowchart in Figure 2 (where we merged Scenario 3 and 4 to a single heterogeneous scenario). Some of the correlations of highly enriched genes in monocyte progenitors seem to originate from heterogeneity (to a lesser extent than genes enriched in neutrophil progenitors but more than the TFs, as can be seen in Figure 5d).

**FIG. E11.**
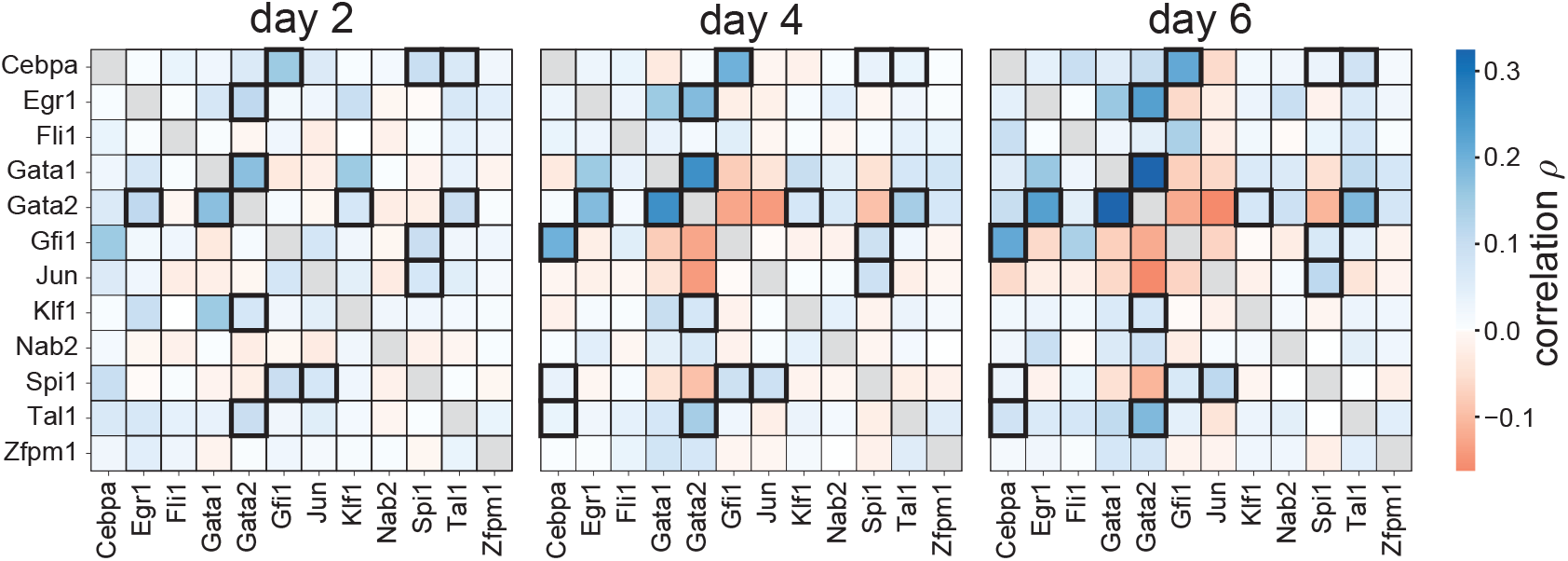
The LARRY dataset is of a dynamic system undergoing differentiation. We plot gene co-expression for the 12 TFs known to drive murine myeloid differentiation at each of the three time points. When we compare these matrices, we find that parts of the GRN are not static. Bold matrix cells denote stable correlations, that is, correlations which are significant and of consistent type across all three time points. We consider only these interactions for further analysis with TwINFER in Figure 5f.

**FIG. E12.**
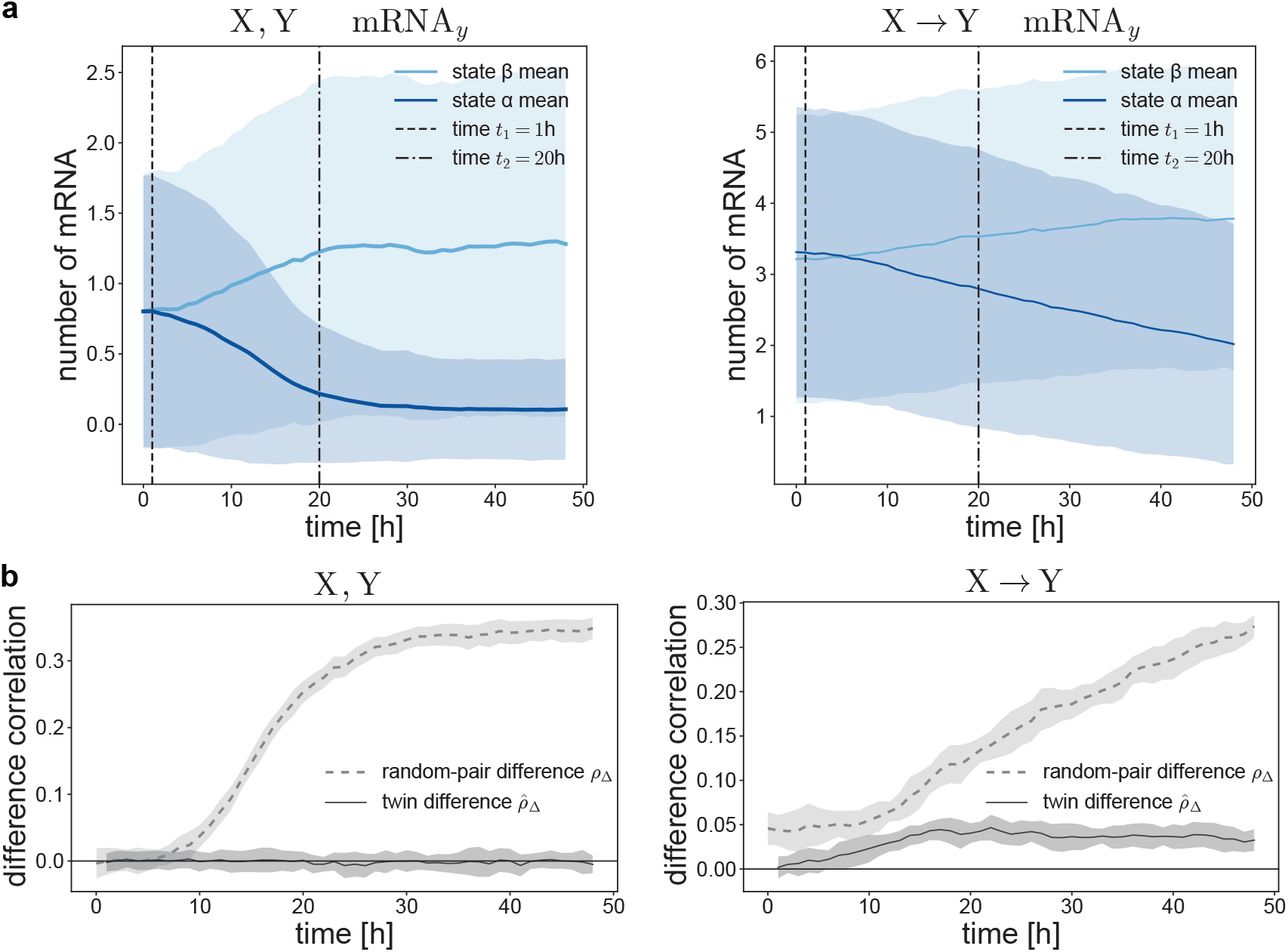
The same scenario as in Figure E2, but now genes X and Y develop their two states after mother cells division, and over time. Mother cells were assigned into two equally sized groups, *α* and *β*, denoting the state of a hidden variable driving the split into two states. The mother cells were otherwise equivalent. They were then allowed to divide, passing their states to their offspring. Therefore, twins share the same state. Once divided, the bursting frequency *k*_on_ of all genes in cells of state *α* (state *β*) slowly drifted towards a lower (higher) value. In **(a)** we plot the mRNA counts over time, for each of the populations. The drift strength was tuned such that between 1 hour (vertical dashed lines) and 20 hours (vertical dot-dashed lines), the target *k*_on_ are reached. This is meant to simulate state differentiation on the time scale of an ideal TwINFER experiment. **(b)** The difference correlations are plotted as a function of time. Random-pair correlation follows the development of the two states. This implies the result of the heterogeneity test will depend on the measurement time.

**FIG. E13.**
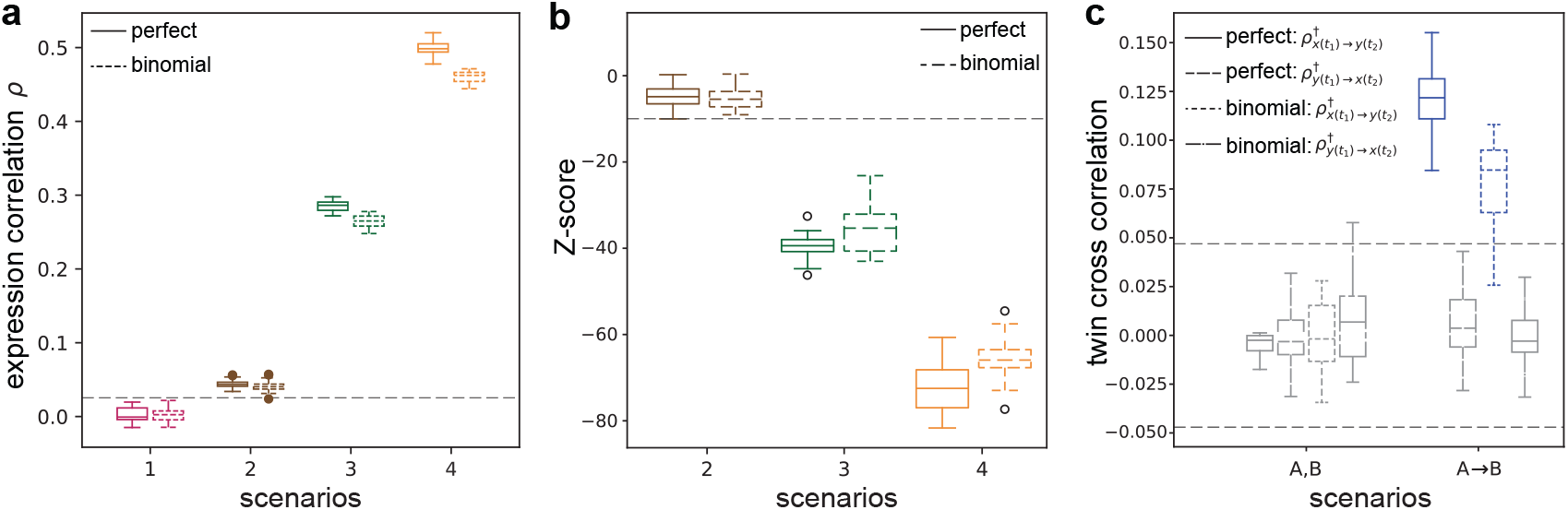
Binomial partitioning of mother cells vs. perfect partitioning. In the main text, we simulated the case where each mother cell partitions its transcriptome and proteome perfectly between the daughter cells. Thus, in perfect partitioning, twins are exactly identical at time zero, with no partition noise. We repeated the simulations with symmetric binomial partitioning, i.e., under the assumption that each transcript and protein has equal probability of joining either daughter cell at cell division (see Ref. [115]). Note that this partition is still symmetric on average, albeit with partition noise. Panels **(a)** and **(b)** correspond to panels (d) and (f) of Figure 2. Panel **(c)** corresponds to the first and second scenarios (from left to right) in panel (b) of Figure 3. In each panel, we show the original results from the main text (perfect partitioning) and the results of repetitions with binomial partitioning. All other simulation parameters remained the same.

