## Supplemental Information for "Leveraging Twin Information for Inference of Gene Regulatory Networks"

### Supplemental Material

#### I. Twin cross-correlations reveal pairwise interaction direction

In order to infer gene-gene interactions in an analytically tractable example, we consider a two-dimensional linear system with directional coupling. Let X and Y be two genes, where the transcript count vector $\mathbf{c}(t) = [x(t), y(t)]^\top$ evolves in time according to Langevin dynamics (time is measured in units of transcript mean degradation time)

$$\frac{d\mathbf{c}}{dt} = -A\mathbf{c}(t) + \eta(t), \tag{S1}$$

where the interaction matrix is

$$A = \begin{pmatrix} 1 & 0 \\ -\lambda & 1 \end{pmatrix}, \tag{S2}$$

and the white Gaussian noise vector is $\eta(t) = [\eta_x(t), \eta_y(t)]^\top$, such that

$$\langle \eta_i(t_1)\eta_j(t_2)\rangle = 2D\delta_{ij}\delta(t_2 - t_1) \quad \forall i, j \in \{x, y\}. \tag{S3}$$

The solution is given by

$$\mathbf{c}(t) = e^{-At}\mathbf{c}_0 + \int_0^t e^{-A(t-s)}\eta(s)\,ds. \tag{S4}$$

We define twin cells as two independent trajectories, $\mathbf{c}(t)$ and $\mathbf{c}'(t)$, initialized at the same value

$$\mathbf{c}(0) = \mathbf{c}'(0) = \mathbf{c}_0 = \begin{pmatrix} x_0 \\ y_0 \end{pmatrix}, \tag{S5}$$

but driven by independent noise realizations

$$\langle \eta_i(t_1)\,\eta'_j(t_2)\rangle = 0 \quad \forall i, j \in \{x, y\}. \tag{S6}$$

From the general solution in Eq. (S4), the cross-covariance matrix between x and y across the population of all twins is the $2X2$ matrix

$$\hat{\mathrm{K}}^\dagger(t_1, t_2) = \begin{pmatrix} \langle x(t_1)x'(t_2)\rangle & \langle x(t_1)y'(t_2)\rangle \\ \langle y(t_1)x'(t_2)\rangle & \langle y(t_1)y'(t_2)\rangle \end{pmatrix} \equiv \begin{pmatrix} \hat{\mathrm{K}}^\dagger_{x(t_1)\to x(t_2)} & \hat{\mathrm{K}}^\dagger_{x(t_1)\to y(t_2)} \\ \hat{\mathrm{K}}^\dagger_{y(t_1)\to x(t_2)} & \hat{\mathrm{K}}^\dagger_{y(t_1)\to y(t_2)} \end{pmatrix} \tag{S7}$$

where the brackets are expectations taken over twin pairs. More explicitly,

$$\hat{\mathrm{K}}^\dagger(t_1, t_2) = \left\langle \left[e^{-At_1}\mathbf{c}_0 + \int_0^{t_1} e^{-A(t_1-s)}\eta(s)\,ds\right]\left[e^{-At_2}\mathbf{c}_0 + \int_0^{t_2} e^{-A(t_2-u)}\eta'(u)\,du\right]^\top \right\rangle, \tag{S8}$$

Expanding, we obtain

$$\begin{aligned}\hat{\mathrm{K}}^\dagger(t_1, t_2) = &\underbrace{e^{-At_1}\mathbf{c}_0\mathbf{c}_0^\top e^{-A^\top t_2}}_{\text{deterministic} \times \text{deterministic}} \underbrace{+ e^{-At_1}\mathbf{c}_0 \left\langle \left(\int_0^{t_2} e^{-A(t_2-u)}\eta'(u)\,du\right)^\top \right\rangle}_{\text{deterministic} \times \text{noise (zero mean)}} \\ &\underbrace{+ \left\langle \int_0^{t_1} e^{-A(t_1-s)}\eta(s)\,ds \right\rangle \mathbf{c}_0^\top e^{-A^\top t_2}}_{\text{noise} \times \text{deterministic (zero mean)}} + \underbrace{\left\langle \left(\int_0^{t_1} e^{-A(t_1-s)}\eta(s)\,ds\right)\left(\int_0^{t_2} e^{-A(t_2-u)}\eta'(u)\,du\right)^\top \right\rangle}_{\text{noise} \times \text{noise}}.\end{aligned} \tag{S9}$$

The second and third terms vanish because $\langle \eta(t) \rangle = 0$. The fourth term also vanishes by use of Eq. (S6). Concluding, the only surviving contribution to the twin cross-covariance comes from the deterministic term

$$\hat{K}^{\dagger}(t_1,t_2) = e^{-At_1}\mathbf{c}_0\mathbf{c}_0^{\top}e^{-A^{\top}t_2}. \tag{S10}$$

Expanding the matrix exponentials, and after some algebra, we obtain

$$e^{-At} = e^{-t}\begin{pmatrix} 1 & 0 \\ \lambda t & 1 \end{pmatrix}, \quad \text{and} \quad e^{-A^{\top}t_2} = e^{-t_2}\begin{pmatrix} 1 & \lambda t_2 \\ 0 & 1 \end{pmatrix}. \tag{S11}$$

Computing the off-diagonal terms in Eq. (S7) we get

$$\hat{K}^{\dagger}(t_1,t_2) = e^{-(t_1+t_2)}\begin{pmatrix} x_0^2 & x_0y_0 + \lambda t_2 x_0^2 \\ x_0y_0 + \lambda t_1 x_0^2 & y_0^2 + \lambda(t_1+t_2)x_0y_0 + \lambda^2 t_1 t_2 x_0^2 \end{pmatrix}. \tag{S12}$$

The asymmetry between the two off-diagonal cross-covariances reflects the directional nature of the regulatory interaction from gene X to gene Y

$$\hat{K}^{\dagger}_{x(t_1)\to y(t_2)} - \hat{K}^{\dagger}_{y(t_1)\to x(t_2)} = \lambda(t_2 - t_1)x_0^2 e^{-(t_2+t_1)}. \tag{S13}$$

Importantly, setting $t_1 = 0$,

$$\hat{K}^{\dagger}_{y(t_1=0)\to x(t_2)} = e^{-t_2}x_0y_0 \tag{S14}$$

is independent of the interaction parameter and decays exponentially, while

$$\hat{K}^{\dagger}_{x(t_1=0)\to y(t_2)} = e^{-t_2}(x_0y_0 + \lambda t_2 x_0^2) \tag{S15}$$

exhibits transient growth on short timescales and the effect is proportional to the interaction parameter $\lambda$. Overall, setting $t_1 = 0$ in Eq. (S12) we obtain

$$\hat{K}^{\dagger}(0,t_2) = e^{-t_2}\begin{pmatrix} x_0^2 & x_0y_0 + \lambda t_2 x_0^2 \\ x_0y_0 & y_0^2 + \lambda t_2 x_0 y_0 \end{pmatrix}. \tag{S16}$$

A more realistic scenario is the case in which different pairs of twins are initialized with $x_0$ and $y_0$ that are drawn from the steady-state distribution of two genes with dynamics governed by Eq. (S1). Specifically, we want the ensemble average of Eq. (S10) with respect to the random initial condition

$$\langle \hat{K}^{\dagger}(t_1,t_2) \rangle = \langle e^{-At_1}\mathbf{c}_0\mathbf{c}_0^{\top}e^{-A^{\top}t_2} \rangle = e^{-At_1}\langle \mathbf{c}_0\mathbf{c}_0^{\top} \rangle e^{-A^{\top}t_2} = e^{-At_1}K_{\text{ss}}e^{-A^{\top}t_2}. \tag{S17}$$

where $K_{\text{ss}}$ is the steady-state of the gene covariance $K(t) = \langle \mathbf{c}(t)\mathbf{c}(t)^{\top} \rangle$. We can find the steady state by solving the Lyapunov equation

$$AK_{\text{ss}} + K_{\text{ss}}A^{\top} = Q, \tag{S18}$$

where $Q = \langle \eta(t)\eta(t)^{\top} \rangle = 2DI$ is the noise covariance matrix. In the case of Eq. (S2), we obtain

$$K_{\text{ss}} = D\begin{pmatrix} 1 & \frac{\lambda}{2} \\ \frac{\lambda}{2} & 1 + \frac{\lambda^2}{2} \end{pmatrix}. \tag{S19}$$

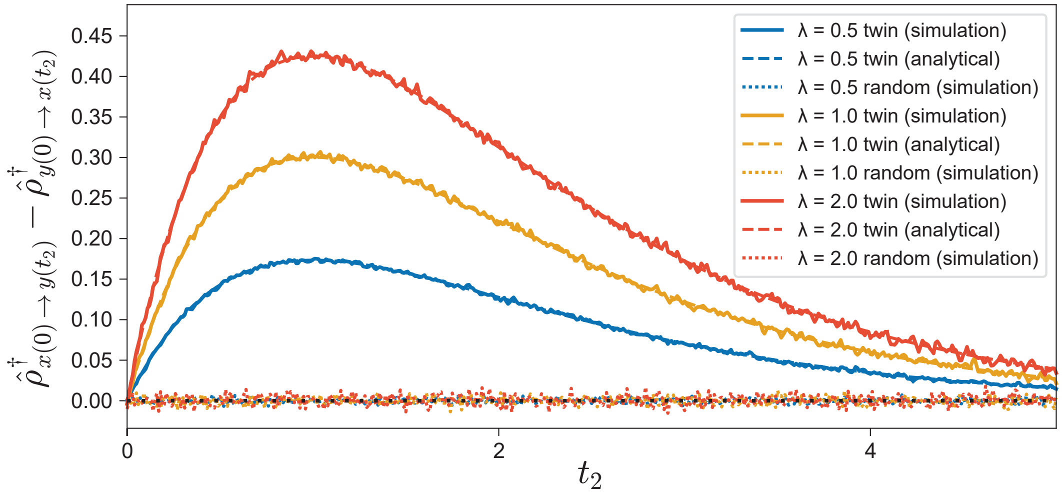


FIG. S1. Twin cross-correlations are asymmetric due to the directional regulation X $\rightarrow$ Y. We plot the difference between the forward and reverse cross-correlations $\hat{\rho}^{\dagger}_{x(t_1)\to y(t_2)} - \hat{\rho}^{\dagger}_{y(t_1)\to x(t_2)}$ as function of the measurement time $t_2$ for various regulatory strengths $\lambda$ ranging from 0.5 to 2.0. We set $t_1 = 0$, $D = 1.25$. Solid lines represent empirical estimates from simulated twin trajectories ($8 \cdot 10^5$ pairs), dashed lines show the analytical solution in Eq. (S24). Twin pairs are initiated from a joint position drawn from the steady-state distribution. Dotted lines show empirical results for the same number of independently initialized (random) pairs.

Plugging Eq. (S19) in Eq. (S17) we get

$$\langle \hat{\mathrm{K}}^{\dagger}(t_1,t_2)\rangle = De^{-(t_1+t_2)} \begin{pmatrix} 1 & \frac{\lambda}{2}(1+2t_2) \\ \frac{\lambda}{2}(1+2t_1) & 1+\frac{\lambda^2}{2}(1+t_1+t_2+2t_1t_2) \end{pmatrix}, \tag{S20}$$

and for the case where $t_1 = 0$

$$\langle \hat{\mathrm{K}}^{\dagger}(0,t_2)\rangle = De^{-t_2} \begin{pmatrix} 1 & \frac{\lambda}{2}(1+2t_2) \\ \frac{\lambda}{2} & 1+\frac{\lambda^2}{2}(1+t_2) \end{pmatrix}, \tag{S21}$$

Finally, we will show that the asymmetry observed in the covariance matrix is carried onto the Pearson correlation matrix where the initial state is drawn from the steady-state distribution. Element-wise, we can define the matrix by

$$\hat{\rho}^{\dagger}_{i(t_1)\to j(t_2)} = \frac{\langle \hat{\mathrm{K}}^{\dagger}_{i(t_1)\to j(t_2)}\rangle}{\sqrt{\mathrm{K}_{\mathrm{ss,ii}}\,\mathrm{K}_{\mathrm{ss,jj}}}} \quad \forall i,j \in \{x,y\}, \tag{S22}$$

where we used the fact that $\langle \hat{\mathrm{K}}^{\dagger}_{i(t)\to i(t)}\rangle = \mathrm{K}_{\mathrm{ss,ii}}$ since the system is assumed to be initiated in steady state. Plugging Eqs. (S19) and (S21) in Eq. (S22), we obtain

$$\hat{\rho}^{\dagger}(0,t_2) = e^{-t_2} \begin{pmatrix} 1 & \frac{\lambda(1+2t_2)}{2\sqrt{1+\frac{\lambda^2}{2}}} \\ \frac{\lambda}{2\sqrt{1+\frac{\lambda^2}{2}}} & 1+\frac{\lambda^2 t_2}{2+\lambda^2} \end{pmatrix}. \tag{S23}$$

The difference between the two off-diagonal cross-correlations is

$$\hat{\rho}^{\dagger}_{x(0)\to y(t_2)} - \hat{\rho}^{\dagger}_{y(0)\to x(t_2)} = \frac{\lambda t_2}{\sqrt{1+\frac{\lambda^2}{2}}} e^{-t_2}. \tag{S24}$$

In Figure S1 we show that the difference between the twin cross-covariances matches Monte-Carlo simulations well. We further observe that twin cross-correlations are directional, whereas random-pair cross-correlations vanish. This is in complete accordance with the behavior observed for the non-linear model in the main text (Figure 3).

**II. Feedback amplifies and prolongs twin cross-correlation**

Let us now consider bidirectional pairwise interactions, replacing the interaction matrix in Eq. (S2) with

$$A = \begin{pmatrix} 1 & -\lambda_{yx} \\ -\lambda_{xy} & 1 \end{pmatrix}, \tag{S25}$$

with $\lambda_{ij}$ being the interaction parameter for gene I regulating gene J. This small exercise will turn out useful in the next section. Solving Eq. (S18) with the new interaction matrix, we obtain

$$\mathrm{K_{ss}} = D \begin{pmatrix} \frac{2-\lambda_{xy}\lambda_{yx}+\lambda_{yx}^2}{2-2\lambda_{xy}\lambda_{yx}} & \frac{\lambda_{xy}+\lambda_{yx}}{2-2\lambda_{xy}\lambda_{yx}} \\ \frac{\lambda_{xy}+\lambda_{yx}}{2-2\lambda_{xy}\lambda_{yx}} & \frac{2+\lambda_{xy}^2-\lambda_{xy}\lambda_{yx}}{2-2\lambda_{xy}\lambda_{yx}} \end{pmatrix}. \tag{S26}$$

We continue the derivation as in the previous section, but the expressions become too cumbersome to restate here. When we set $t_1 = 0$ and $\lambda_{xy} = \lambda_{yx} = \lambda$, the expressions simplify to

$$\hat{\rho}^\dagger(0,t_2) = e^{-t_2} \begin{pmatrix} \cosh(\lambda t_2) + \lambda \sinh(\lambda t_2) & \lambda \cosh(\lambda t_2) + \sinh(\lambda t_2) \\ \lambda \cosh(\lambda t_2) + \sinh(\lambda t_2) & \cosh(\lambda t_2) + \lambda \sinh(\lambda t_2) \end{pmatrix}. \tag{S27}$$

As expected from the symmetry of the system, the off-diagonal cross-correlations are equal in this case. More importantly, they are non-zero, and the bidirectional case can be detected by computing the cross-correlations in both directions. When $\lambda_{xy} \neq \lambda_{yx}$, the matrix becomes asymmetric in a way that reflects the relative strength of the two interactions.

As shown in Figure S2, the cross-correlation magnitude is greater in the bidirectional regulation than in the unidirectional regulation. In the unidirectional case (X $\rightarrow$ Y), the cross-correlations from X to Y scale with $\lambda$ (in accordance with Eq. (S1)). In bidirectional regulation (X $\rightleftarrows$ Y), there is a positive feedback between the two genes, which increases the shared component between them, resulting in a larger cross-correlation.

Moreover, bidirectional interaction has a slower decay rate. To see this, consider the eigenvalues of the interaction matrix. For unidirectional regulation, both eigenvalues are equal to 1, so the correlation decays as a single exponential $e^{-t_2}$, which does not depend on the value of $\lambda$. On the other hand, bidirectional interaction has eigenvalues $1 \pm \lambda$,

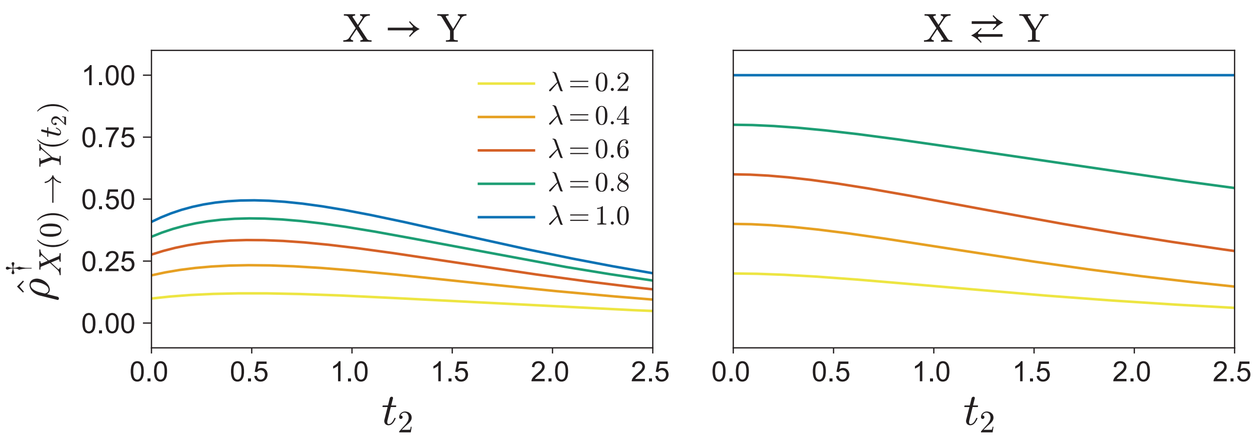


FIG. S2. Twin cross-correlations vs. the measurement time $t_2$. We set $t_1 = 0$. Left: unidirectional regulation (X $\rightarrow$ Y), given by Eq. (S23). Right: bidirectional regulation ($X \rightleftarrows Y$), given by Eq. (S27). Curves correspond to increasing regulatory strength $\lambda_{xy} = \lambda_{yx} = \lambda$ within the stable regime ($|\lambda| \leq 1$). The positive feedback between the two genes results in substantially higher cross-correlations.

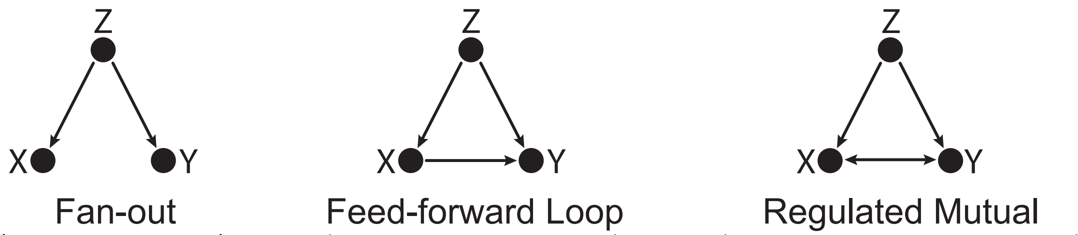


$$\begin{pmatrix} 1 & 0 & -\lambda_{zx} \\ 0 & 1 & -\lambda_{zy} \\ 0 & 0 & 1 \end{pmatrix} \quad \begin{pmatrix} 1 & 0 & -\lambda_{zx} \\ -\lambda_{xy} & 1 & -\lambda_{zy} \\ 0 & 0 & 1 \end{pmatrix} \quad \begin{pmatrix} 1 & -\lambda_{yx} & -\lambda_{zx} \\ -\lambda_{xy} & 1 & -\lambda_{zy} \\ 0 & 0 & 1 \end{pmatrix}$$

FIG. S3. Triplet GRN motifs and their corresponding interaction matrix $A$. From left to right: fan-out, feed-forward loop, and regulated mutual. When using correlation-based inference methods, and when the ground truth is fan-out or feed-forward loop, it is often the case that the X–Y edge of regulated mutual is a false positive due to Z acting as a confounding factor.

producing two exponential terms $e^{-(1-\lambda)t_2}$ and $e^{-(1+\lambda)t_2}$. As $\lambda$ increases within the stable regime $|\lambda| \leq 1$, the term $1-\lambda$ becomes smaller, yielding a slower decay of the correlation.

Lastly, note that the linear amplification effect does not carry over to the non-linear model, see Figure 3 and discussion in the main text.

#### III. Twin correlations distinguish between fan-out, feed-forward loop and regulated mutual motifs

We now expand the two-gene system of Sections I and II by adding a gene Z that can pairwise interact with genes X and Y. For simplicity, let us assume that only activation is possible. For each edge in the triplet, there are four options: no interaction, unidirectional interaction in the forward direction, unidirectional interaction in the reverse direction, and bidirectional interaction. Overall, the addition of a single gene increases the number of possible configurations from 4 to 64.

Some of these configurations are known to be difficult to infer using correlation-based inference schemes. Among these, the fan-out motif (Figure S3, left) is especially notorious, as it is also an ubiquitous motif in real GRN, along with the feed-forward loop [33]. These two important motifs are indistinguishable based on single-timepoint correlations.

When the ground truth motif is fan-out, correlation-based inference schemes often infer it as the regulated mutual motif (Figure S3, right). In fact, the fan-out motif represents a classical problem in inference, where gene Z acts as a confounding factor. Moreover, also the feed-forward loop motif (Figure S3, center) can result in the regulated mutual being inferred. Therefore, when the result of a correlation-based inference method is a regulated mutual, it is likely that at least one of the inferred interactions X $\rightarrow$ Y and X $\leftarrow$ Y is a false positive. Given the ubiquity of fan-out motifs in GRNs we face the question: Upon inferring a regulated mutual motif, can we leverage twin information to assess the likelihood of this edge being a false positive?

First, let us show that regular steady-state gene co-expression (single sample) cannot inform us. The generalization of Eq. (S1) is straightforward. We consider $\mathbf{c}(t) = [x(t), y(t), z(t)]^\top$, $\eta(t) = [\eta_x(t), \eta_y(t), \eta_z(t)]^\top$ and the motif-corresponding matrix $A$ in Figure S3. We then use Eq. (S18) again. Solving, we obtain for the fan-out case

$$\mathrm{K}_{\mathrm{ss}} = D \begin{pmatrix} 1+\frac{\lambda_{zx}^2}{2} & \frac{\lambda_{zx}\lambda_{zy}}{2} & \frac{\lambda_{zx}}{2} \\ \frac{\lambda_{zx}\lambda_{zy}}{2} & 1+\frac{\lambda_{zy}^2}{2} & \frac{\lambda_{zy}}{2} \\ \frac{\lambda_{zx}}{2} & \frac{\lambda_{zy}}{2} & 1 \end{pmatrix}, \tag{S28}$$

and the corresponding Pearson correlations

$$\rho = \begin{pmatrix} 1 & \frac{\lambda_{zx}\lambda_{zy}}{\sqrt{(2+\lambda_{zx}^2)(2+\lambda_{zy}^2)}} & \frac{\lambda_{zx}}{\sqrt{2(2+\lambda_{zx}^2)}} \\ \frac{\lambda_{zx}\lambda_{zy}}{\sqrt{(2+\lambda_{zx}^2)(2+\lambda_{zy}^2)}} & 1 & \frac{\lambda_{zy}}{\sqrt{2(2+\lambda_{zy}^2)}} \\ \frac{\lambda_{zx}}{\sqrt{2(2+\lambda_{zx}^2)}} & \frac{\lambda_{zy}}{\sqrt{2(2+\lambda_{zy}^2)}} & 1 \end{pmatrix}. \tag{S29}$$

For simplicity, we set $t_1 = 0$ and $\lambda_{ij} = \lambda$ for all the interactions. The off-diagonal gene-expression correlations are simplified to

$$\begin{aligned} \rho_{xy} &= \rho_{yx} = \frac{\lambda^2}{2+\lambda^2}, \\ \rho_{xz} &= \rho_{zx} = \rho_{yz} = \rho_{zy} = \frac{\lambda}{\sqrt{2(2+\lambda^2)}}. \end{aligned} \tag{S30}$$

Comparing these expressions, the order of the correlations reverses at $\lambda = \sqrt{2}$: for $0 < \lambda < \sqrt{2}$ we have $\rho_{xz} = \rho_{yz} > \rho_{xy}$, while for $\lambda > \sqrt{2}$ we obtain $\rho_{xz} = \rho_{yz} < \rho_{xy}$.

Similarly, for the feed-forward loop we obtain

$$\mathrm{K_{ss}} = D \begin{pmatrix} 1+\frac{\lambda_{zx}^2}{2} & \frac{1}{8}\left[\lambda_{xy}(4+3\lambda_{zx}^2)+4\lambda_{zx}\lambda_{zy}\right] & \frac{\lambda_{zx}}{2} \\ \frac{1}{8}\left[\lambda_{xy}(4+3\lambda_{zx}^2)+4\lambda_{zx}\lambda_{zy}\right] & \frac{1}{8}\left[8+\lambda_{xy}^2(4+3\lambda_{zx}^2)+6\lambda_{xy}\lambda_{zx}\lambda_{zy}+4\lambda_{zy}^2\right] & \frac{1}{4}\left(\lambda_{xy}\lambda_{zx}+2\lambda_{zy}\right) \\ \frac{\lambda_{zx}}{2} & \frac{1}{4}\left(\lambda_{xy}\lambda_{zx}+2\lambda_{zy}\right) & 1 \end{pmatrix}, \tag{S31}$$

and the gene co-expression matrix can be written element-wise as

$$\begin{aligned} \rho_{xy} &= \rho_{yx} = \frac{\frac{1}{8}\left[\lambda_{xy}(4+3\lambda_{zx}^2)+4\lambda_{zx}\lambda_{zy}\right]}{\sqrt{\left(1+\frac{\lambda_{zx}^2}{2}\right)\frac{1}{8}\left[8+\lambda_{xy}^2(4+3\lambda_{zx}^2)+6\lambda_{xy}\lambda_{zx}\lambda_{zy}+4\lambda_{zy}^2\right]}}, \\ \rho_{xz} &= \rho_{zx} = \frac{\frac{\lambda_{zx}}{2}}{\sqrt{1+\frac{\lambda_{zx}^2}{2}}}, \\ \rho_{yz} &= \rho_{zy} = \frac{\frac{1}{4}\left(\lambda_{xy}\lambda_{zx}+2\lambda_{zy}\right)}{\sqrt{\frac{1}{8}\left[8+\lambda_{xy}^2(4+3\lambda_{zx}^2)+6\lambda_{xy}\lambda_{zx}\lambda_{zy}+4\lambda_{zy}^2\right]}}, \\ \rho_{xx} &= \rho_{yy} = \rho_{zz} = 1. \end{aligned} \tag{S32}$$

Setting $\lambda_{ij} = \lambda$, we get

$$\begin{aligned} \rho_{xy} &= \rho_{yx} = \frac{\lambda(3\lambda^2+4\lambda+4)}{2\sqrt{3\lambda^6+6\lambda^5+14\lambda^4+12\lambda^3+24\lambda^2+16}}, \\ \rho_{xz} &= \rho_{zx} = \frac{\lambda}{\sqrt{2(\lambda^2+2)}}, \\ \rho_{yz} &= \rho_{zy} = \frac{\lambda(\lambda+2)}{\sqrt{2(3\lambda^4+6\lambda^3+8\lambda^2+8)}}, \\ \rho_{xx} &= \rho_{yy} = \rho_{zz} = 1. \end{aligned} \tag{S33}$$

For all $\lambda > 0$, $\rho_{xy}$ is always the largest. The only change in ranking occurs between $\rho_{xz}$ and $\rho_{yz}$ at $\lambda \approx 1.151$,

$$\begin{aligned} \rho_{xy} > &\rho_{yx} > \rho_{xz}, && 0 < \lambda < 1.151, \\ \rho_{xy} > &\rho_{xz} = \rho_{yz}, && \lambda \approx 1.151, \\ \rho_{xy} > &\rho_{xz} > \rho_{yz}, && \lambda > 1.151. \end{aligned} \tag{S34}$$

The regulated mutual terms are given by:

$$
\begin{aligned}
&\mathrm{K}_{\mathrm{ss},xx} = D\frac{(-4+\lambda_{xy}\lambda_{yx})(-2+(\lambda_{xy}-\lambda_{yx})\lambda_{yx}-\lambda_{zx}^2)+6\lambda_{yx}\lambda_{zx}\lambda_{zy}+3\lambda_{yx}^2\lambda_{zy}^2}{2(-4+\lambda_{xy}\lambda_{yx})(-1+\lambda_{xy}\lambda_{yx})},\\
&\mathrm{K}_{\mathrm{ss},xy} = \mathrm{K}_{\mathrm{ss},yx} = D\frac{4\lambda_{yx}-\lambda_{xy}\left(-4+\lambda_{yx}(\lambda_{xy}+\lambda_{yx})-3\lambda_{zx}^2\right)+2(2+\lambda_{xy}\lambda_{yx})\lambda_{zx}\lambda_{zy}+3\lambda_{yx}\lambda_{zy}^2}{2(-4+\lambda_{xy}\lambda_{yx})(-1+\lambda_{xy}\lambda_{yx})},\\
&\mathrm{K}_{\mathrm{ss},xz} = \mathrm{K}_{\mathrm{ss},zx} = D\frac{2\lambda_{zx}+\lambda_{yx}\lambda_{zy}}{4-\lambda_{xy}\lambda_{yx}},\\
&\mathrm{K}_{\mathrm{ss},yy} = D\frac{8-\lambda_{xy}^3\lambda_{yx}+\lambda_{xy}^2\left(4+\lambda_{yx}^2+3\lambda_{zx}^2\right)+4\lambda_{zy}^2-\lambda_{xy}\left(-6\lambda_{zx}\lambda_{zy}+\lambda_{yx}(6+\lambda_{zy}^2)\right)}{2(-4+\lambda_{xy}\lambda_{yx})(-1+\lambda_{xy}\lambda_{yx})},\\
&\mathrm{K}_{\mathrm{ss},yz} = \mathrm{K}_{\mathrm{ss},zy} = D\frac{\lambda_{xy}\lambda_{zx}+2\lambda_{zy}}{4-\lambda_{xy}\lambda_{yx}},\\
&\mathrm{K}_{\mathrm{ss},zz} = D.
\end{aligned}
\tag{S35}
$$

Using covariances above in Eq. (S35), gene-expression correlations are computed as:

$$
\begin{aligned}
&\rho_{xy} = \rho_{yx} = \frac{N_{xy}}{\sqrt{N_{xx}N_{yy}}},\\
&\rho_{xz} = \rho_{zx} = \frac{\left(2\lambda_{zx}+\lambda_{yx}\lambda_{zy}\right)\sqrt{2(-4+\lambda_{xy}\lambda_{yx})(-1+\lambda_{xy}\lambda_{yx})}}{\left(4-\lambda_{xy}\lambda_{yx}\right)\sqrt{N_{xx}}},\\
&\rho_{yz} = \rho_{zy} = \frac{\left(\lambda_{xy}\lambda_{zx}+2\lambda_{zy}\right)\sqrt{2(-4+\lambda_{xy}\lambda_{yx})(-1+\lambda_{xy}\lambda_{yx})}}{\left(4-\lambda_{xy}\lambda_{yx}\right)\sqrt{N_{yy}}},\\
&\rho_{xx} = \rho_{yy} = \rho_{zz} = 1.
\end{aligned}
\tag{S36}
$$

where $N$ is defined as:

$$
\begin{aligned}
N_{xx} &= (-4+\lambda_{xy}\lambda_{yx})\left(-2+(\lambda_{xy}-\lambda_{yx})\lambda_{yx}-\lambda_{zx}^2\right)+6\lambda_{yx}\lambda_{zx}\lambda_{zy}+3\lambda_{yx}^2\lambda_{zy}^2,\\
N_{xy} &= 4\lambda_{yx}-\lambda_{xy}\left(-4+\lambda_{yx}(\lambda_{xy}+\lambda_{yx})-3\lambda_{zx}^2\right)+2(2+\lambda_{xy}\lambda_{yx})\lambda_{zx}\lambda_{zy}+3\lambda_{yx}\lambda_{zy}^2,\\
N_{yy} &= 8-\lambda_{xy}^3\lambda_{yx}+\lambda_{xy}^2\left(4+\lambda_{yx}^2+3\lambda_{zx}^2\right)+4\lambda_{zy}^2-\lambda_{xy}\left(-6\lambda_{zx}\lambda_{zy}+\lambda_{yx}(6+\lambda_{zy}^2)\right).
\end{aligned}
$$

With all regulatory strengths equal to $\lambda$, we get

$$
\begin{aligned}
&\rho_{xy} = \rho_{yx} = \frac{\lambda(\lambda^2+2)}{\lambda^3+\lambda^2-\lambda+2},\\
&\rho_{xz} = \rho_{zx} = \rho_{yz} = \rho_{zy} = -\frac{\lambda}{\lambda-2}\sqrt{\frac{\lambda^3-2\lambda^2-\lambda+2}{\lambda^3+\lambda^2-\lambda+2}},\\
&\rho_{xx} = \rho_{yy} = \rho_{zz} = 1.
\end{aligned}
\tag{S37}
$$

For the regulated mutual motif, the stability of the linear system requires all eigenvalues of the interaction matrix $A$ to have a positive real part. In the symmetric case ($\lambda_{xy} = \lambda_{yx} = \lambda_{zx} = \lambda_{zy} = \lambda$), the X $\leftrightarrow$ Y submatrix (upper left $2\times 2$ matrix in Figure S3) has eigenvalues $1\pm\lambda$, and thus the steady state exists only when $1-\lambda > 0$, i.e., $0 < \lambda < 1$. Within this stable regime we always find $\rho_{xy} > \rho_{xz} = \rho_{yz}$.

The first thing to note is that all three covariance matrices contain a non-negative $\rho_{xy} = \rho_{yx}$, even though there is no interaction between X and Y in the fan-out case. Indeed, Z is a confounding variable. Not knowing the strengths of the interactions, we cannot rely on $\rho_{xy}$ to infer the existence or non-existence of an interaction between X and Y. In this sense, Pearson correlations suffer from the same limitation as the raw covariances.

We thus need a more sophisticated measure. We again turn to twin information and consider the cross-correlations defined in Section I. We can compute the covariances for the three triplets by plugging in Eq. (S17), the corresponding $A$ matrices in Figure S3 and the $\mathrm{K}_{\mathrm{ss}}$ matrix in Eqs. (S28)-(S35). For simplicity, we set $t_1 = 0$ and all $\lambda_{ij} = \lambda$ for all the interactions. In the fan-out case, after some algebra, we obtain

$$\langle \hat{\mathrm{K}}^{\dagger}(0,t_2)\rangle = De^{-t_2}\begin{pmatrix} \frac{1}{2}\left[2+\lambda^2(1+t_2)\right] & \frac{1}{2}\lambda^2(1+t_2) & \frac{1}{2}\lambda \\ \frac{1}{2}\lambda^2(1+t_2) & \frac{1}{2}\left[2+\lambda^2(1+t_2)\right] & \frac{1}{2}\lambda \\ \frac{1}{2}\lambda(1+2t_2) & \frac{1}{2}\lambda(1+2t_2) & 1 \end{pmatrix}. \tag{S38}$$

Similarly, for the feed-forward loop we obtain

$$\langle \hat{\mathrm{K}}^{\dagger}(0,t_2)\rangle = De^{-t_2}\times \tag{S39}$$
$$\begin{pmatrix} \frac{1}{2}\left(2+\lambda^2(1+t_2)\right) & \frac{1}{8}\lambda\left(4+4\lambda+3\lambda^2+4(2+\lambda+\lambda^2)t_2+2\lambda^2t_2^2\right) & \frac{1}{2}\lambda \\ \frac{1}{8}\lambda\left(4+4\lambda+3\lambda^2+2\lambda(2+\lambda)t_2\right) & \frac{1}{8}\left(8+8\lambda^2+6\lambda^3+3\lambda^4+\lambda^2(8+6\lambda+3\lambda^2)t_2+\lambda^3(2+\lambda)t_2^2\right) & \frac{1}{4}\lambda(2+\lambda) \\ \frac{1}{2}\lambda(1+2t_2) & \frac{1}{4}\lambda(2+\lambda+2(2+\lambda)t_2+2\lambda t_2^2) & 1 \end{pmatrix},$$

and for the regulated mutual

$$\langle \hat{\mathrm{K}}^{\dagger}(0,t_2)\rangle = De^{-t_2}\begin{pmatrix} \frac{(2+\lambda^2)\cosh(\lambda t_2)+\lambda(-1+\lambda^2+3\sinh(\lambda t_2))}{(-2+\lambda)(-1+\lambda^2)} & \frac{-\lambda+\lambda^3+3\lambda\cosh(\lambda t_2)+(2+\lambda^2)\sinh(\lambda t_2)}{(-2+\lambda)(-1+\lambda^2)} & \frac{\lambda}{2-\lambda} \\ \frac{-\lambda+\lambda^3+3\lambda\cosh(\lambda t_2)+(2+\lambda^2)\sinh(\lambda t_2)}{(-2+\lambda)(-1+\lambda^2)} & \frac{(2+\lambda^2)\cosh(\lambda t_2)+\lambda(-1+\lambda^2+3\sinh(\lambda t_2))}{(-2+\lambda)(-1+\lambda^2)} & \frac{\lambda}{2-\lambda} \\ \frac{\lambda(1+3\sinh(\lambda t_2)+(2+\lambda)\cosh(\lambda t_2))}{(-2+\lambda)} & \frac{\lambda(1+3\sinh(\lambda t_2)+(2+\lambda)\cosh(\lambda t_2))}{(-2+\lambda)} & 1 \end{pmatrix}. \tag{S40}$$

Plugging these covariances in Eq. (S22), we obtain the Pearson correlations corresponding to the fan-out case

$$\hat{\rho}^{\dagger}(0,t_2) = e^{-t_2}\begin{pmatrix} \frac{2+\lambda^2+\lambda^2t_2}{2+\lambda^2} & \frac{\lambda^2(1+t_2)}{2+\lambda^2} & \frac{\lambda}{\sqrt{2(2+\lambda^2)}} \\ \frac{\lambda^2(1+t_2)}{2+\lambda^2} & \frac{2+\lambda^2+\lambda^2t_2}{2+\lambda^2} & \frac{\lambda}{\sqrt{2(2+\lambda^2)}} \\ \frac{\lambda(1+2t_2)}{\sqrt{2(2+\lambda^2)}} & \frac{\lambda(1+2t_2)}{\sqrt{2(2+\lambda^2)}} & 1 \end{pmatrix}. \tag{S41}$$

Similarly, for the feed-forward loop

$$\hat{\rho}^{\dagger}(0,t_2) = e^{-t_2}\begin{pmatrix} \frac{2+\lambda^2+\lambda^2t_2}{2+\lambda^2} & \frac{\lambda\left(4+\lambda(4+3\lambda)+4(2+\lambda+\lambda^2)t_2+2\lambda^2t_2^2\right)}{2\sqrt{(2+\lambda^2)(8+\lambda^2(8+3\lambda(2+\lambda)))}} & \frac{\lambda}{\sqrt{2(2+\lambda^2)}} \\ \frac{\lambda(4+\lambda(4+3\lambda)+2\lambda(2+\lambda)t_2)}{2\sqrt{(2+\lambda^2)(8+\lambda^2(8+3\lambda(2+\lambda)))}} & \frac{8+\lambda^2(8+3\lambda(2+\lambda))+\lambda^2t_2(8+3\lambda(2+\lambda)+\lambda(2+\lambda)t_2)}{8+\lambda^2(8+3\lambda(2+\lambda))} & \frac{\lambda(2+\lambda)}{\sqrt{2(8+\lambda^2(8+3\lambda(2+\lambda)))}} \\ \frac{\lambda(1+2t_2)}{\sqrt{2(2+\lambda^2)}} & \frac{\lambda(2+\lambda+2t_2(2+\lambda+\lambda t_2))}{\sqrt{2(8+\lambda^2(8+3\lambda(2+\lambda)))}} & 1 \end{pmatrix}, \tag{S42}$$

and for the regulated mutual (in the stable regime $\lambda \leq 1$)

$$\begin{aligned}
\hat{\rho}^{\dagger}_{x(0)\to x(t_2)} &= \frac{e^{-t_2}\left[(2+\lambda^2)\cosh(\lambda t_2)+\lambda(-1+\lambda^2+3\sinh(\lambda t_2))\right]}{2+\lambda(-1+\lambda+\lambda^2)}, \\
\hat{\rho}^{\dagger}_{x(0)\to y(t_2)} &= \frac{e^{-t_2}\left[-\lambda+\lambda^3+3\lambda\cosh(\lambda t_2)+(2+\lambda^2)\sinh(\lambda t_2)\right]}{2+\lambda(-1+\lambda+\lambda^2)}, \\
\hat{\rho}^{\dagger}_{x(0)\to z(t_2)} &= -\frac{e^{-t_2}\lambda}{(-2+\lambda)\sqrt{\frac{2+\lambda(-1+\lambda+\lambda^2)}{(-2+\lambda)(-1+\lambda)(1+\lambda)}}}, \\
\hat{\rho}^{\dagger}_{y(0)\to x(t_2)} &= \frac{e^{-t_2}\left[-\lambda+\lambda^3+3\lambda\cosh(\lambda t_2)+(2+\lambda^2)\sinh(\lambda t_2)\right]}{2+\lambda(-1+\lambda+\lambda^2)}, \\
\hat{\rho}^{\dagger}_{y(0)\to y(t_2)} &= \frac{e^{-t_2}\left[(2+\lambda^2)\cosh(\lambda t_2)+\lambda(-1+\lambda^2+3\sinh(\lambda t_2))\right]}{2+\lambda(-1+\lambda+\lambda^2)},
\end{aligned} \tag{S43}$$

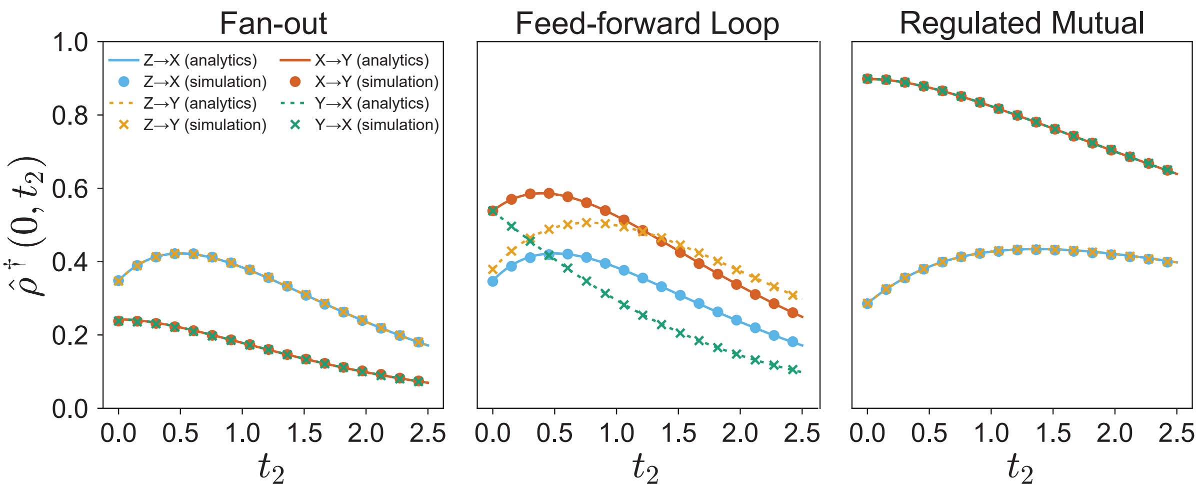


FIG. S4. Twin cross-correlations distinguish between the three triplets from Figure S3. For each triplet, we plot the cross-correlation $\hat{\rho}^{\dagger}_{i(0)\to j(t_2)}$ vs. $t_2$, for $ij \in \{xy, yx, xz, yz\}$. We set $t_1 = 0$ and $\lambda_{ij} = \lambda = 0.8$ for all $ij$, and $D = 1.25$. The analytical curves are given by Eqs. (S41)-(S43). Monte Carlo simulations of $3 \cdot 10^5$ pairs of twins cells fit the analytical expression well. The order is given by Eq. (S44). Importantly, the order is unique for each motif, allowing inference of the X–Y edge (which is indistinguishable with regular gene-expression correlations $\rho$). Note that $10^4$ pairs are enough for the identification of the triplets, but slightly noisier, here we use more pairs to demonstrate the excellent agreement with theory.

$$
\begin{aligned}
\hat{\rho}^{\dagger}_{y(0)\to z(t_2)} &= -\frac{e^{-t_2}\lambda}{(-2+\lambda)\sqrt{\frac{2+\lambda(-1+\lambda+\lambda^2)}{(-2+\lambda)(-1+\lambda)(1+\lambda)}}},\\
\hat{\rho}^{\dagger}_{z(0)\to x(t_2)} &= -\frac{e^{-t_2}\left(-2+2e^{\lambda t_2}+\lambda\right)}{(-2+\lambda)\sqrt{\frac{2+\lambda(-1+\lambda+\lambda^2)}{(-2+\lambda)(-1+\lambda)(1+\lambda)}}},\\
\hat{\rho}^{\dagger}_{z(0)\to y(t_2)} &= -\frac{e^{-t_2}\left(-2+2e^{\lambda t_2}+\lambda\right)}{(-2+\lambda)\sqrt{\frac{2+\lambda(-1+\lambda+\lambda^2)}{(-2+\lambda)(-1+\lambda)(1+\lambda)}}},\\
\hat{\rho}^{\dagger}_{z(0)\to z(t_2)} &= e^{-t_2}.
\end{aligned}
$$

One can appreciate that, while we can compute the correlation matrix exactly for any network, the expressions quickly become extremely bulky. The Pearson correlation matrices were given here for completeness, after setting $t_1 = 0$ and all $\lambda_{i,j} = \lambda$, but we can set any parameter in the full bulkier equations not shown here. The qualitative behavior does not change.

Inspecting the resulting cross-correlation matrices

$$\hat{\rho}^{\dagger}_{zy} = \hat{\rho}^{\dagger}_{zx} > \underline{\hat{\rho}^{\dagger}_{xy}} = \underline{\hat{\rho}^{\dagger}_{yx}}, \quad \text{fan-out (for all } \lambda \text{ when } t_2 > 1/\sqrt{2}), \tag{S44a}$$

$$\hat{\rho}^{\dagger}_{zy} > \underline{\hat{\rho}^{\dagger}_{xy}} > \underline{\hat{\rho}^{\dagger}_{yx}} > \hat{\rho}^{\dagger}_{zx}, \quad \text{feed-forward loop}, \tag{S44b}$$

$$\underline{\hat{\rho}^{\dagger}_{xy}} = \underline{\hat{\rho}^{\dagger}_{yx}} > \hat{\rho}^{\dagger}_{zx} = \hat{\rho}^{\dagger}_{zy}, \quad \text{regulated mutual (only stable when } \lambda \le 1). \tag{S44c}$$

Thus, each of the triplets has a different order of cross-correlations, enabling us to distinguish between the different motifs, as demonstrated in Figure S4. Due to symmetry, some of the correlations share the same rank. If we treat genes that share a rank as a single rank group, one can show that allowing for different values $\lambda_{i,j}$ does not change the order (within each group, equalities can become inequalities). Note that for $t_2 < 1/\sqrt{2}$, there exists a value of $\lambda_c(t_2) = \sqrt{2(1+2t_2)^2/(1-2t_2^2)}$ above which the fan-out order is reversed. As $t_2 \to 0$ we have $\lambda_c(t_2) \to \sqrt{2}$. Recall that time is measured here in units of the mean transcript degradation time, and so to ensure a unique order for each motif regardless of the interaction strength, we require that $t_2$ exceeds $1/\sqrt{2}$ times the mean degradation time (a few hours).

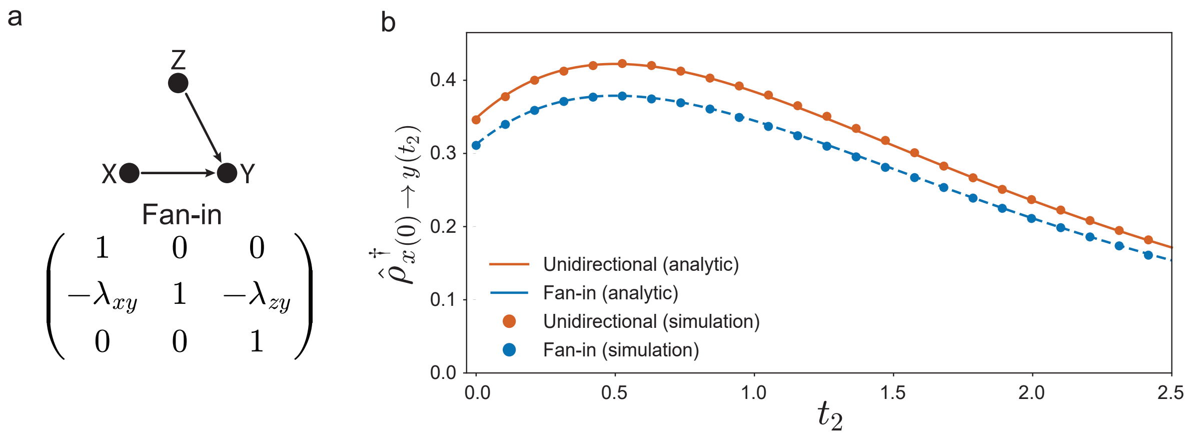


FIG. S5. **(a)** The fan-in motif and its interaction matrix. **(b)** A comparison between the twin cross-correlations $\hat{\rho}^{\dagger}_{x(0)\to y(t_2)}$ of the fan-in motif (solid orange line) vs the two-gene unidirectional regulation (dashed blue line), plotted against the measurement time $t_2$. We set $t_1 = 0$, $\lambda_{xy} = \lambda_{zy} = 0.8$, $D = 1.25$. The circles denote Monte-Carlo simulation results of $3 \cdot 10^5$ twin pairs.

We have just established that cross-correlations are sufficient to distinguish between the triplets in the linear model. However, in the non-linear model, the order of the cross-correlations in the regulated mutual is similar to that of fan-out. This is accompanied by a considerable attenuation of all correlations. In the main text and in Figure E9, we have systematically narrowed down the source of the attenuation to a sub-motif of the regulated mutual—the fan-in motif (Figure S5a). In the main text we present an alternative route for inference of the regulated mutual. Nonetheless, for completeness, let us explain where the models differ.

First, we establish that fan-in attenuates correlations in the linear model as well. Indeed, by repeating the procedure used for the other three motifs, we obtain for fan-in

$$\mathrm{K}_{ss} = D\begin{pmatrix} 1 & \frac{\lambda_{xy}}{2} & 0 \\ \frac{\lambda_{xy}}{2} & 1+\frac{\lambda_{xy}^2+\lambda_{zy}^2}{2} & \frac{\lambda_{zy}}{2} \\ 0 & \frac{\lambda_{zy}}{2} & 1 \end{pmatrix}, \tag{S45}$$

and the corresponding cross-correlations

$$\begin{aligned} \hat{\rho}^{\dagger}_{x(0)\to y(t)} &= \frac{\lambda_{xy}\left(t+\frac{1}{2}\right)e^{-t}}{\sqrt{1+\frac{\lambda_{xy}^2+\lambda_{zy}^2}{2}}}, \qquad & \hat{\rho}^{\dagger}_{z(0)\to y(t)} &= \frac{\lambda_{zy}\left(t+\frac{1}{2}\right)e^{-t}}{\sqrt{1+\frac{\lambda_{xy}^2+\lambda_{zy}^2}{2}}} \\ \hat{\rho}^{\dagger}_{y(0)\to x(t)} &= \frac{\frac{\lambda_{xy}}{2}e^{-t}}{\sqrt{1+\frac{\lambda_{xy}^2+\lambda_{zy}^2}{2}}} & \hat{\rho}^{\dagger}_{y(0)\to z(t)} &= \frac{\frac{\lambda_{zy}}{2}e^{-t}}{\sqrt{1+\frac{\lambda_{xy}^2+\lambda_{zy}^2}{2}}} \\ \hat{\rho}^{\dagger}_{x(0)\to z(t)} &= \hat{\rho}^{\dagger}_{z(0)\to x(t)} = 0. \end{aligned} \tag{S46}$$

In Figure S5b we compare the correlation of X $\to$ Y between the fan-in and the unidirectional two-gene system. We observe a slight reduction in the correlation due to the fan-in motif. It seems, therefore, that all of the phenomena observed in the linear model carry over but one: the mutual activation feedback loop. Recall that in Section II, we saw that positive feedback between two genes in a bidirectional interaction amplifies cross-correlations in the linear regime. Indeed, the regulated mutual contains the sub-motif X $\leftrightarrow$ Y, resulting in a dramatic amplification. In fact, the cross-correlation between X and Y is the highest.

To conclude, the positive feedback enhances correlations in the linear analysis, albeit not in the non-linear model. Hence, regulated mutual, which embeds a feedback motif, behaves differently. While motif identification is straightforward in the linear model and relies solely on cross-correlations, in the non-linear model we also invoke difference correlations to fully distinguish the triplets (see the main text and Figure 4).
